# Temperature-induced methylome changes during asexual reproduction trigger transcriptomic and phenotypic changes in *Fragaria vesca*

**DOI:** 10.1101/2022.10.09.511489

**Authors:** YuPeng Zhang, Guangxun Fan, Tuomas Toivainen, Torstein Tengs, Igor Yakovlev, Paal Krokene, Timo Hytönen, Carl Gunnar Fossdal, Paul E. Grini

**Affiliations:** EVOGENE, Department of Biosciences, University of Oslo, 0313 Oslo, Norway; Department of Molecular Plant Biology, Norwegian Institute of Bioeconomy Research, 1431 Ås, Norway; Department of Agricultural Sciences, Viikki Plant Science Centre, University of Helsinki, 00014 Helsinki, Finland

## Abstract

Plants must quickly adapt to a changing environment in order to maintain their fitness. One rapid adaptation mechanism that promotes plasticity is epigenetic memory, which may provide long-lived organisms the precious time needed to adapt to climate change. In this study, we used the perennial plant *Fragaria vesca* as a model to determine how the methylome and transcriptome adapt to elevated temperatures (28 vs. 18 °C) over three asexual generations. Changes in flowering time, stolon number, and petiole length were induced in responses to temperature treatment in one or more ecotypes after three asexual generations in a manner indicative of an epigenetic memory. Induced methylome changes differed between four different ecotypes from Norway, Iceland, Italy, and Spain, but there were also some shared responses. Elevated temperature conditions induced significant phenotypic and methylation changes, particularly in the Norwegian ecotype. Most of the differentially methylated regions (DMRs) were in the CHG context, and most CHG and CHH DMRs were hypermethylated. Notably, the four ecotypes shared only eight CHG DMR peaks. Several differentially methylated genes (DMGs) also showed a change in gene expression. Ecotype-specific methylation and expression patterns were observed for genes related to gibberellin metabolism, flowering time, epigenetics. Furthermore, when repetitive elements (REs) were found near (±2 kb) or inside a gene, they showed a negative correlation with gene expression. In conclusion, phenotypic changes induced by elevated temperatures during asexual reproduction were accompanied by changes in DNA methylation patterns. Also, positional influences of REs impacted gene expression, indicating that DNA methylation may be involved in both general and ecotype-specific phenotypic plasticity in *F. vesca*.

## Introduction

Climate change is a great challenge for the biosphere and exerts a strong selection pressure, pushing organisms to rapidly adapt to a changing environment. Plants growing in extreme climates are under particularly strong natural selection to synchronize their growth with the environment. In addition to allelic variation, epigenetic modifications provide a mechanism for short-term adaptation to environmental change and preservation of genetic variation. By increasing genetic plasticity epigenetic modifications may influence many aspects of plant phenology, including flowering time (Basu 2020; Agustí *et al.* 2020; Forestan *et al.* 2020), flower color (Jiang, Zhang & Ma 2020; Sobral, Neylan, Narbona & Dirzo 2021), vegetative growth (Li, Hu & Chu 2021a), kernel color, and vernalization (Bastow *et al.* 2004; Csorba, Questa, Sun & Dean 2014). Epigenetic mechanisms, such as DNA methylation, histone modification, and small RNA-dependent pathways, modify the accessibility of DNA in chromatin and thus modulate gene expression and alter the plant phenotype. In the context of environmental change, direct changes in DNA methylation due to environmental impacts can be triggered by changes in various biotic or abiotic parameters, including increased temperature (Bräutigam *et al.* 2013; Wibowo *et al.* 2016; Liu *et al.* 2021). Canonical DNA methylation refers to 5-methylcytosine, where cytosine bases of the DNA are modified by the addition of a methyl group (Stein, Sciaky-Gallili, Razin & Cedar 1983). The three canonical DNA methylation contexts are CGN, CHG, and CHH (where N can be any base and H can be A, C or T). CGN is the most common context, but the latter two contexts are also common in plants, while they are rare in mammals and other organisms (Pélissier, Thalmeir, Kempe, Sänger & Wassenegger 1999; Law & Jacobsen 2010). Plants have three main classes of methyltransferases which transfer and covalently attach methyl groups onto DNA (Jullien, Susaki, Yelagandula, Higashiyama & Berger 2012; Tirot, Bonnet & Jullien 2022) The METHYLTRANSFERASE 1 (MET1), CHROMOMETHYLASE 3 (CMT3) and DOMAINS REARRANGED METHYLTRANSFERASE 1 and 2 (DRM1/DRM2) are unique to plants and have different roles in the DNA methylation pathways (Lindroth *et al.* 2001; Cao *et al.* 2003; Pontes *et al.* 2006). MET1 methylates cytosines in the CGN context, CMT3 is in charge of CHG methylation, while DRM is known to play a role both in maintaining methylation marks trough replication and as a *de novo* DNA methylation methyltransferase (Lindroth *et al.* 2001; Cao *et al.* 2003; Pontes *et al.* 2006).

The RNA-directed DNA methylation pathway (RdDM) is unique to plants. RdDM DNA methylation occurs genome-wide, but less frequently in pericentromeric heterochromatin (Schoft *et al.* 2009; Matzke & Mosher 2014). RdDM is induced by small RNAs through two RNA polymerases, RNA POLYMERASE IV (Pol IV) and RNA POLYMERASE V (Pol V) (Haag & Pikaard 2011; Eun *et al.* 2012). Canonical RdDM needs Pol IV with RNA DEPENDENT RNA POLYMERASE 2 (RDR2) to synthesize double-stranded RNA which is then cut by DICER LIKE 3 (DCL3) into 24 nt small interfering RNAs (siRNAs). These siRNAs bind to ARGONAUTE 4 (AGO4) and Pol V to recruit DRM2 methyltransferases that methylate target DNA (Pikaard, Haag, Pontes, Blevins & Cocklin 2012; Ji & Chen 2012).

Epigenetic regulators can also act in concert. For example, the concerted action of DNA methyltransferases CHROMOMETHYLASE 2 (CMT2), CMT3, DRM1, and DRM2 play a very important role in histone 3 lysine 9 (H3K9) methylation in the chromatin. The KRYPTONITE (KYP) protein reads both CHG and CHH DNA methylation contexts via its SRA domains and then performs histone methylation on nearby chromatin (Jackson, Lindroth, Cao & Jacobsen 2002; Jackson *et al.* 2004; Malagnac, Bartee & Bender 2002; Johnson *et al.* 2007). Such CHG and CHH DNA methylation depends on CMT2 and CMT3 as well as KYP and its homologs SUPPRESSOR OF VARIEGATION 3-9 HOMOLOGs 5 (SUVH5) and SUVH6 (Jackson *et al.* 2002; Johnson *et al.* 2007). The histone demethylases LYSINE-SPECIFIC HISTONE DEMETHYLASE 1 HOMOLOG 1 (LDL1) and LDL2 remove dimethylation and trimethylation from histone 3 lysine 4 (H3K4) to initiate siRNA biogenesis by Pol IV (Greenberg *et al.* 2013). RELATIVE OF EARLY FLOWERING 6 (REF6), a Jumonji domain-containing histone demethylase, recognizes a specific CHG-containing motif in hypomethylated regions, thus causing targeted histone demethylation (Li *et al.* 2016; Qiu *et al.* 2019).

Both abiotic and biotic environmental factors have been shown to affect DNA methylation in plants. When *Arabidopsis thaliana* faces high osmotic stress, an epigenetic memory is induced that can be passed onto the next generation, mainly by female gametes (Wibowo *et al.* 2016). Inorganic phosphate (Pi) starvation in rice (*Oryza sativa*) induces CHH hypermethylation in transposable elements which targets Pi-starvation-induced (PSI) genes (Secco *et al.* 2015). Not only stress-related epigenetic memories can be inherited by the next generation. In toadflax (*Linaria vulgaris*), natural epimutation of *Lcyc*, a homolog of the cycloidea gene, changes flower symmetry (Gustafsson 1979; Cubas, Vincent & Coen 1999). When an *A. thaliana* lineage with a hypomethylated genome is crossed with a normal wild type, massive methylation and expression changes occur in the next generation (Rigal *et al.* 2016). Moreover, differentially methylated regions (DMRs) segregate as distinct loci in recombinant inbred lines (RILs) in both soybean (*Glycine max*) and *A. thaliana* (Johannes *et al.* 2009; Nery, Diers, Xu, Stacey & Ecker 2013). More intriguingly, organ-specific methylation markers are inherited by asexual offspring and can be maintained even through the following meiosis (Wibowo *et al.* 2018).

In the conifer Norway spruce (*Picea abies*), increased temperature sums during zygotic and somatic embryogenesis induces an epigenetic memory. This memory affects the timing of bud burst and bud set in genetically identical progenies (epitypes) in a predictable and reproducible manner, and the effect appears to be life-lasting (Yakovlev, Fossdal & Johnsen 2010; Yakovlev, Lee, Rotter & Olsen 2014; Yakovlev, Carneros, Lee, Olsen & Fossdal 2016; Carneros, Yakovlev, Viejo, Olsen & Fossdal 2017). These Norway spruce epitypes differed significantly in their transcriptome and small RNA profile (Yakovlev *et al.* 2010, 2014). Following embryogenesis at epitype-inducing temperature conditions, 329 out of 735 putative epigenetic-related gene orthologs were differentially expressed (Yakovlev *et al.* 2016). Besides epigenetic-related genes, gene expression analysis also showed that most DEHYDRINS (DHNs), EARLY BUD BREAK 1 (EBB1), and FLOWERING LOCUS T LIKE 2 (FTL2) genes were differentially expressed between epitypes both at and before bud burst in a manner previously reported for ecotype differences (Carneros *et al.* 2017).

We chose *Fragaria vesca (2n = 2x = 14)* as a model to investigate how a perennial plant responds to climate change, and specifically if a temperature-induced epigenetic memory like that observed in Norway spruce also exists in other species. *F. vesca*, also known as woodland strawberry, is common throughout Eurasia. It is one of the ancestors of commercial strawberry and is used as a reference to understand the genomics of the octoploid commercial strawberry (Shulaev *et al.* 2008; Liston, Cronn & Ashman 2014). *F. vesca* has two reproduction systems: sexual reproduction with production of achenes and asexual reproduction through stolon formation. Because both reproductive systems have a relatively short generation cycle, *F. vesca* is a potent system for studying climatic responses in perennial plants. In this study, we tested for lasting phenotypic impacts of different temperature conditions (28°C and 18°C) experienced during one to three generations of asexual propagation using four *F. vesca* ecotypes. We quantified DNA methylation in leaf tissues, identified differentially methylated regions (DMRs) induced by the temperature treatment, and compared differentially methylated genes (DMGs), repetitive elements (REs), and other genic features associated with DMRs and differentially expressed genes (DEGs). We demonstrate that *F. vesca* has a temperature-induced epigenetic memory and that phenotypic changes induced by elevated temperatures are accompanied by changes in DNA methylation patterns.

## Materials and Methods

### Plant material and experimental conditions

*Fragaria vesca* ecotypes (genotypes) ‘ES12’ (43.5339ºN, 6.5271ºW, 138 m, Spain), ‘ICE2’ (63.9988ºN, 19.9604ºW, 99 m, Iceland), ‘IT4’ (46.2398ºN, 11.2360ºE, 949 m, Italy), and ‘NOR2’ (69.9395ºN, 23.0964ºE, 23 m, Norway) (Fig 1A) were grown in a growth chamber equipped with Valoya AP67 LED lamps (200 μmol m^−2^s^−1^ photosynthetically active radiation (PAR) at 40-45% humidity). A long day (LD) length of 16h/8h (light/dark) was used for plant vegetative growth and a short day (SD) length of 12h/12h was used to induce flowering. Plants were grown in 400 ml pots and fertilized regularly with a fertilizer solution (N-P-K: 17-4-25, Kekkilä, Finland). Up to three asexual generations were generated at normal temperatures (18 ºC) and at elevated temperatures (28 ºC) to induce epigenetic changes (Fig S1). For the first asexual generation (AS1) stolons from mother plants of the four ecotypes listed above were rooted at both temperatures. Subsequent asexual generations (AS2 and AS3) were generated with 3-week intervals using stolons from previous asexual generations. Two weeks after the initiation of AS1 and AS3, at the time point of 6 hours after lights were turned on, 20 mg of young unfolded leaf tissues was collected from 10 plants per treatment per ecotype for DNA/RNA isolation. Three weeks later, plants were moved to SD conditions at 18 ºC for 5 weeks (AS1) or 6 weeks (AS3) to induce flowering. After SD treatment, petiole length of the youngest fully developed leaf was recorded, and the plants were moved to a common environment (16/8h LD, 18-20 ºC) in the greenhouse for further phenotypic observations. Flowering time was recorded as days to the first open flower, and the number of stolons was recorded every 2 weeks for 14 weeks. AS2 plants were not phenotyped or sampled for molecular analyses.

**Fig 1.**
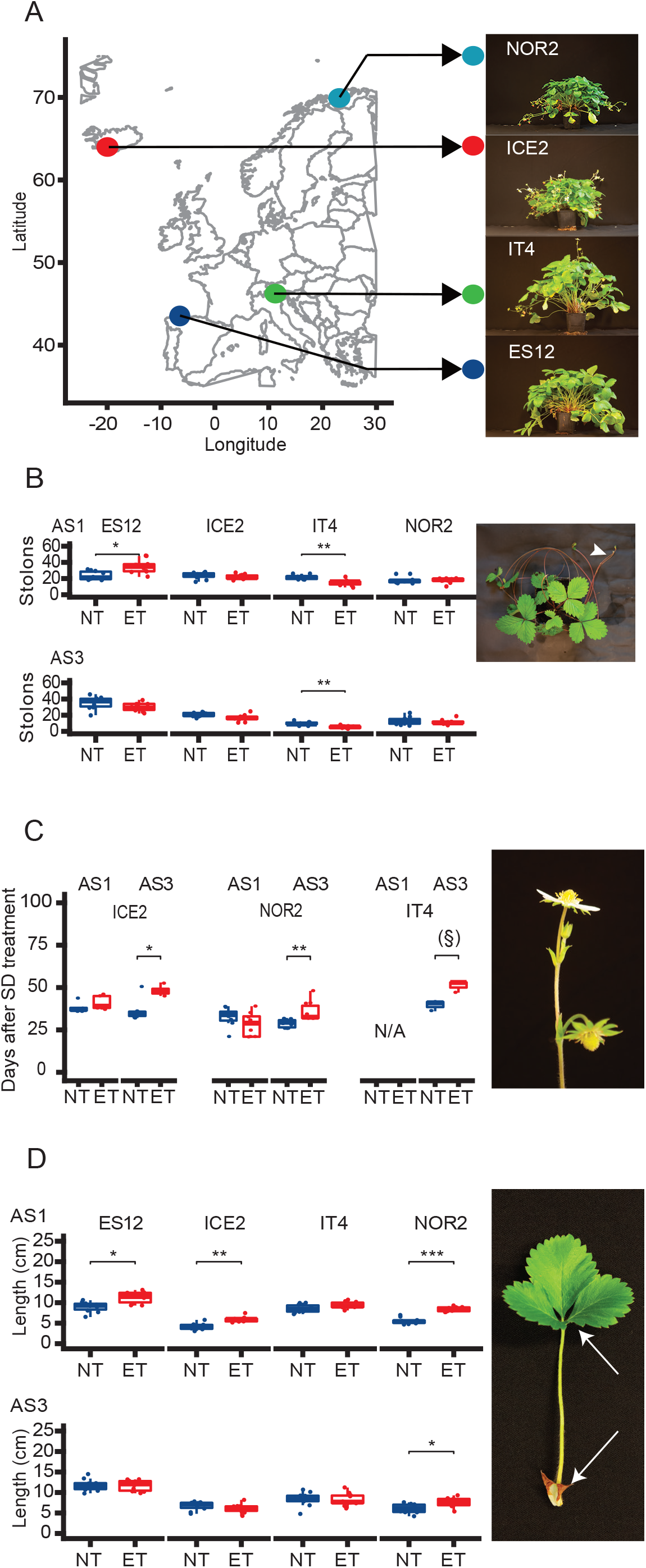
Phenotypic changes in *Fragaria* vesca plants propagated at different temperatures for up to three generations. **(A)** Origin of the four ecotypes used in this study. Total number of stolon **(B)**, flowering time **(C)**, and petiole length **(D)** in the ES12, ICE2, IT4, and NOR2 ecotypes. Plants were propagated under normal (18 ºC) or elevated temperatures (28 ºC) and phenotypes were scored after flower-inducing short-day (SD) treatment. Flowering time was measured as days from the start of SD treatment until the first flower had opened completely. Flowering time could not be determined in ES12 plants since these could not be induced to flower. Arrowhead in images indicates stolon, arrow indicates petiole length. AS1: asexual generation 1; AS3: asexual generation 3. Brackets and asterisks indicate significant differences calculated from Wilcoxon tests: * 0.01 ≤ p < 0.05; ** 0.001 ≤ p < 0.01; *** 0.0001 ≤ p < 0.001. § p=0.06. 95% confidence intervals were used as error bars, average value, 25 quantile and 75 quantile is also shown.

### DNA and RNA isolation

Simultaneous extraction of both genomic DNA and total RNA from leaves was performed as described previously (Zeng, Raffaello, Liu & Asiegbu 2018).

### Bisulfite and RNA-sequencing

Bisulfite library preparation and sequencing were done by BGI using the HiSeq platform. Transcriptome libraries and RNA-seq were also constructed and performed by BGI using the DNB-seq platform. Three independent replicates of bisulfite reads were mapped and analyzed using the default settings in the methylpy pipeline (Schultz *et al.* 2016). Principal component analysis and other analyses of global DNA methylation patterns and methylation of different ecotypes and temperature epitypes were done using methylpy and R with a window size of 50 kb (Schultz *et al.* 2016). Global methylation circos plots were made using shinyCircos with methylation data from the global methylation analysis (Yu, Ouyang & Yao 2018). Plots of gene densities and methylation levels were made using the R package Rideogram (Hao *et al.* 2020). Statistical comparisons of methylation of different genomic features were done using the Wilcoxon test in R. Differentially methylated regions (DMRs) of different methylation contexts were identified using the DMRfind function of methylpy with the following parameters: window size of 250 bp, minimal read coverage of 30, and p-value < 0.003 (Schultz *et al.* 2016). The methylation distribution pattern of genes and repetitive elements were plotted using the R package methimpute (Taudt *et al.* 2018). Body regions of genes and repetitive elements were sectioned into 20 pieces to calculate the methylation degree, whereas flanking regions (±2 kb) were sectioned into 20 pieces of 100 bp each. DMRs were annotated to genomic features using the R package ChIPseeker (Yu, Wang & He 2015). DMRs circos plots were made using shinyCircos (Yu *et al.* 2018). Venn plots were generated using an online tool (https://bioinformatics.psb.ugent.be/webtools/Venn/). Gene ontology (GO) term enrichment analysis was done using the R package clusterprofiler (Wu *et al.* 2021). Any positional effects of repetitive elements were analyzed using the games_howell_test from the R package rstatix. Heatmaps were made using the R package pheatmap.

Three independent replicates of RNA-seq reads were analyzed using the default settings in CLC Genomics Workbench (Qiagen Ltd). Volcano plots were made using the ggplot2 package in R, based on log_2_FoldChange-values and log_10_ p-values from the CLC output. For statistical testing of overlapping DMGs and differentially expressed genes (DEGs) an online tool running Fisher’s exact test was used (http://nemates.org/MA/progs/overlap_stats.html). Repeatable elements (REs, DATASHEET1) were predicted using REPEATMODELER2 with default settings and pseudogenes (DATASHEET2) were predicted using pseudopipe (Zhang *et al.* 2006; Flynn *et al.* 2020). Linear regression analysis was done in R to explore relationships between DNA methylation and genomic features (model: methylation∼gene density + RE density + pseudogene density).

### Accession Numbers

All sequences generated in this study have been deposited in the National Center for Biotechnology Information Sequence Read Archive (https://www.ncbi.nlm.nih.gov/sra) with project number PRJNA879428. Supplemental Data and Supplemental Tables are available at GitHub (https://github.com/sherlock0088/FvAex).

## Results

### Temperature memory induces phenotypic changes

We tested whether a substantial difference in temperature (18 vs. 28 ºC) during asexual propagation would generate heritable epigenetic alterations across three asexual generations in four European *F. vesca* ecotypes (Fig 1A). To uncover any long-term phenotypic changes indicative of temperature-induced epigenetic memory effects, we examined flowering time, stolon number, and petiole length under common garden-conditions after one and three asexual generations (AS1, AS3).

The ES12 ecotype produced significantly more stolons after propagation at 28 ºC than at 18 ºC in AS1 but not in AS3 (Table S1). The IT4 ecotype, on the other hand, produced significantly fewer stolons at 28 ºC both in AS1 and AS3 (Fig 1B, Table S1, Wilcoxon test, 0.05 > p > 0.001). Notably, IT4 plants propagated at 28 ºC stopped generating stolons from week 11 onwards in AS3 (Fig S2, Table S1). Although treatment variation was observed also in the ICE2 and NOR2 ecotypes (Fig S2, Table S1), stolon production was not altered significantly by temperature in these ecotypes.

Temperature treatments affected flowering time considerably (Fig 1C, Table S1). In AS1, only ICE2 and NOR2 could be induced to flower, but there were no significant effects of temperature on flowering time. In AS3, however, flowering occurred significantly later in ICE2 and NOR2 plants propagated at 28 ºC than at 18 ºC (Fig 1C, Table S1, Wilcoxon test, 0.05 > p > 0.001). Three out of ten IT4 replicates flowered in AS3, but there were no significant differences between treatments (Fig 1C, Table S1). The ES12 ecotype did not flower at all under our experimental conditions, indicating that it has different environmental requirements for flowering (possibly shorter photoperiod or cooler temperatures).

Petiole length was significantly longer in AS1 in ES12, ICE2, and NOR2 plants propagated at 28 ºC than at 18 ºC (Fig. 1D, Table S1; Wilcoxon test, 0.05 > p > 0.0001). Petiole length differences disappeared in AS3 for ES12 and ICE2 but persisted for NOR2 (Fig 1D, Table S1, 0.05 > p > 0.0001). In summary, we observed statistically significant differences between temperature conditions for all traits investigated under common garden conditions, indicating the presence of a memory effect of the temperature experienced during asexual propagation.

### Global differences in DNA methylation between ecotypes

We sequenced bisulfite-treated genomic DNA samples from young leaves of all four *F. vesca* ecotypes propagated at 18 and 28 ºC (Fig S1). Quality control showed an overall coverage of >20× for most replicates and comparable mapping rates across ecotypes (Table S2).

First, we compared global cytosine methylation in the CGN, CHG, and CHH methylation contexts at 18 ºC using AS3 plants and found the highest methylation level in the symmetric CGN context (Fig 2A). Out of 8 million CGN sites identified in the *F. vesca* genome, 55-59% were methylated in the different ecotypes (Table S2). In comparison, the average methylation level of CHG and CHH sites was approximately 30% and 6%, respectively (Table S2). The most densely methylated regions on the different chromosomes corresponded with centromeric and pericentromeric heterochromatin (Fig S4) (Qu *et al.* 2017). Although individual *F. vesca* chromosomes showed distinct methylation patterns, these patterns were similar in all four ecotypes (Fig 2A, Fig S3, Fig S4). Nevertheless, at 18 ºC there were some methylation differences between ecotypes (Fig 2B, Fig S5, Table S3). ICE2 was overall hypermethylated compared to all other ecotypes at both a single-chromosome and whole-genome level (Fig 2B, Fig S5, Table S3). We also analyzed the importance of specific trinucleotide contexts of methylated CGN, CHG, and CHH sites on a chromosomal scale. Notably, in the CHG context CCG was less prone to methylation than CTG and CAG in distinct chromosomal regions (Fig S6, S7, S8), as has been observed previously in both monocots and dicots (Gouil & Baulcombe 2016).

**Fig 2.**
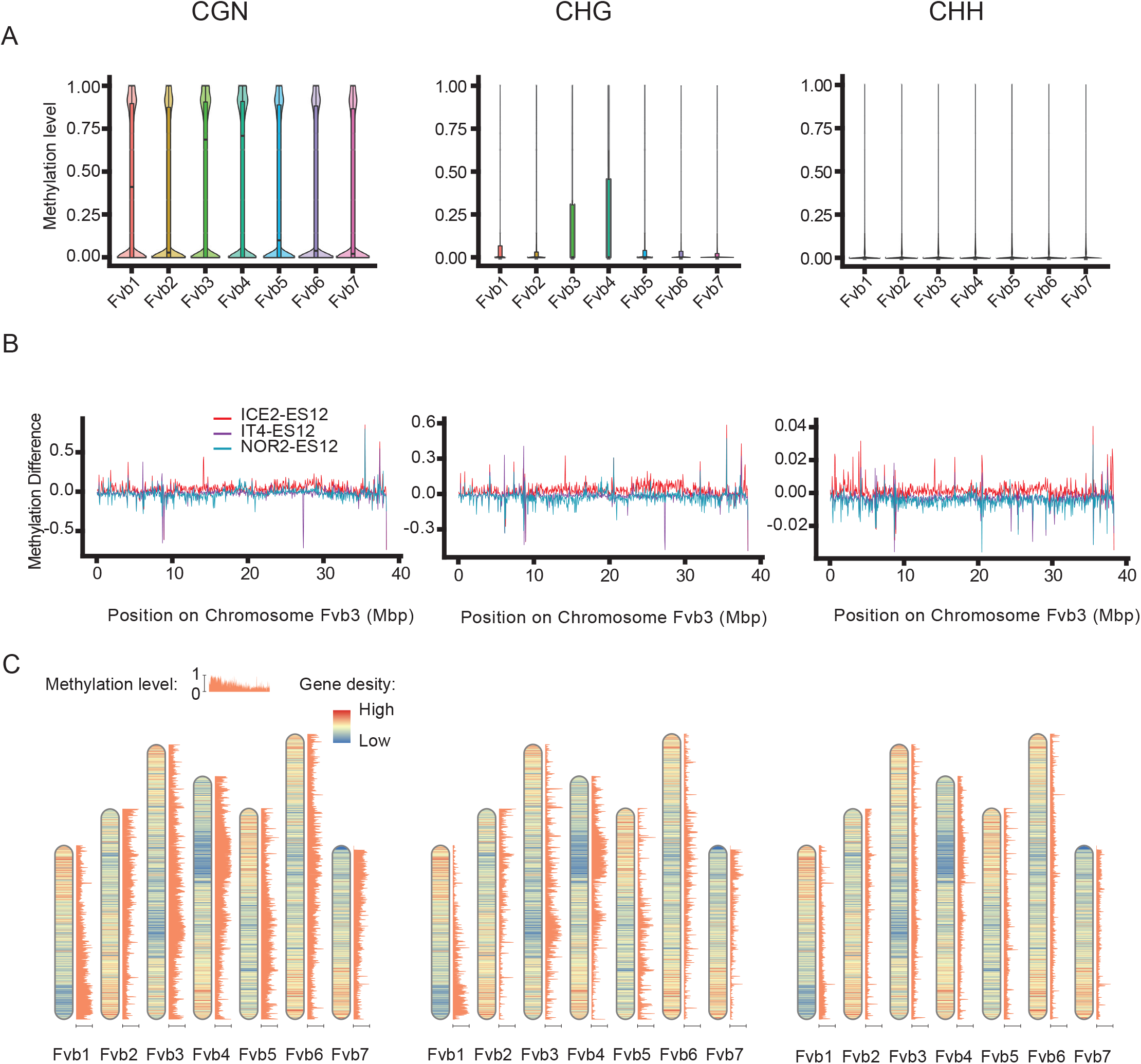
DNA methylation landscape in leaves of *Fragaria vesca* plants propagated at normal temperature conditions (18 ºC). Average methylation values (methylated reads/total reads) were calculated based on the methylation level of each methylated site, discarding any unmethylated sites.**(A)** Violin plots showing the DNA methylation level of different methylation contexts on all seven *F. vesca* (Fv) chromosomes. The 25 quantile and 75 quantile values are indicated by the lower and upper margins of the boxplot and the median is indicated by the black line between quantiles. Error bars show 95% confidence intervals. Ecotype shown here is NOR2. **(B)** Methylation differences between *F. vesca* ecotypes ES12 (Spain), ICE2 (Iceland), IT4 (Italy), and NOR2 (Norway) along chromosome Fvb3. CGN, CHG and CHH methylation patterns of 50 kb windows were calculated along the chromosome using ES12 as a reference. **(C)** Correlation between methylation level and gene density along all seven chromosomes, using a 50 kb window. Ecotype shown here is NOR2.

Linear regression analysis revealed an inverse relationship between protein-coding gene density and the degree of methylation on each chromosome. In all ecotypes, genes were frequently located in low-methylation areas of the chromosomes, regardless of the methylation context (Fig 2C, Fig S9). However, we found a positive correlation to hypermethylated areas for repetitive elements (REs) and pseudogenes (Table S4). Only ∼50% of CGN methylation, ∼55% of CHG methylation, and ∼30% of CHH methylation could be explained by a linear regression model taking all investigated genomic features into account (Table S4).

### Global methylation changes are induced by temperature

Principal Component Analysis (PCA) of methylation profiles in plants propagated at 18 and 28 ºC demonstrated little variability between biological replicates (Fig. 3A). The analysis further highlighted differences in methylation patterns between ecotypes propagated at 18 ºC (Fig 3A). These methylation patterns shifted when plants were propagated at 28 ºC, particularly in the CHG and CHH contexts (Fig 3A). Elevated temperatures changed these two methylation contexts in similar ways across ecotypes and replicates, as changes in methylation followed the same vector directions in the PCA plots. Among the ecotypes, NOR2 had the largest difference between methylation values at 18 and 28 ºC (Fig 3A, Table S5).

**Fig 3.**
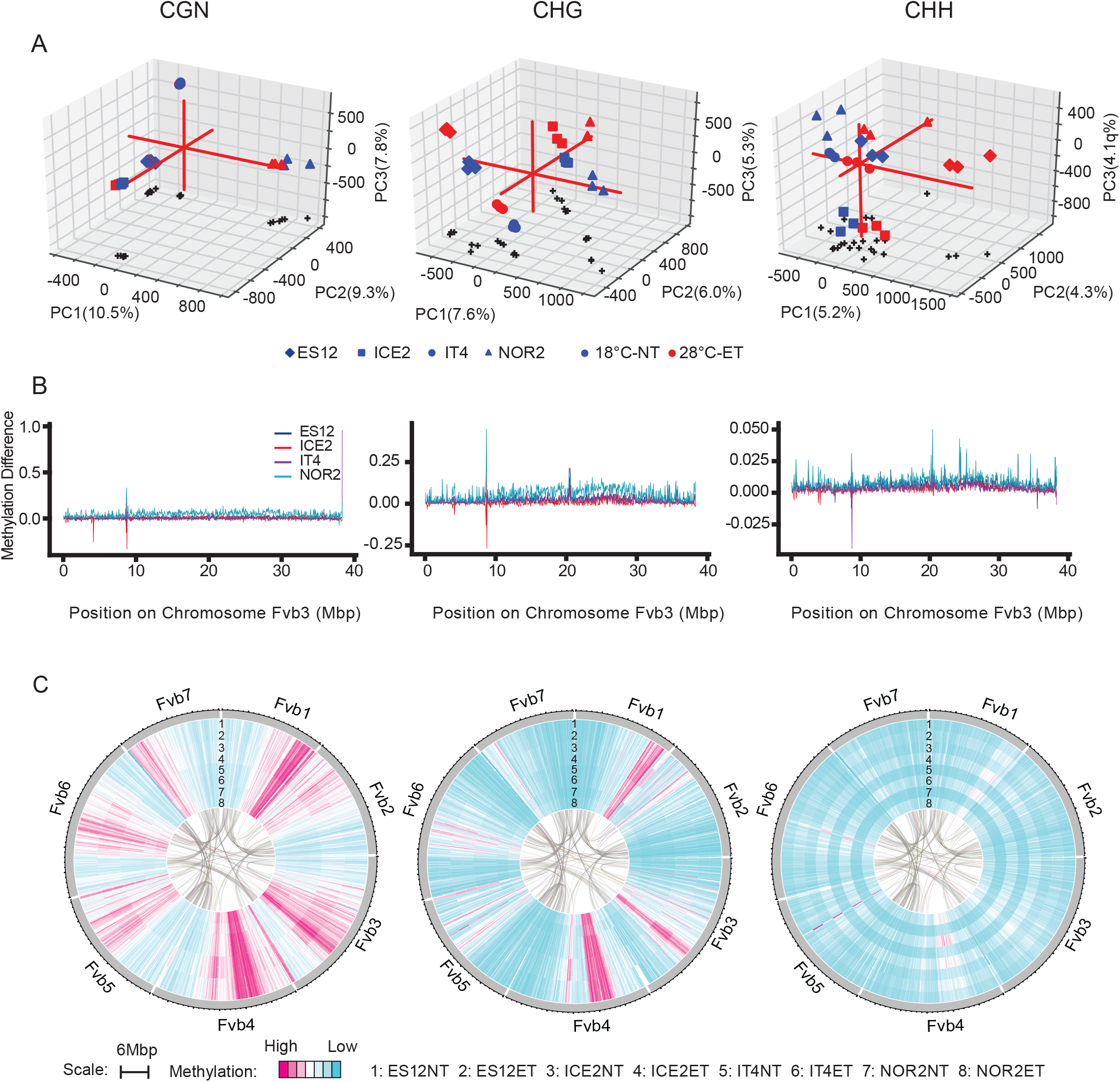
DNA methylation landscape in leaves of four *Fragaria vesca* ecotypes propagated at normal (18 ºC) and elevated (28 ºC) temperature conditions. Ecotypes are from Spain (ES12), Iceland (ICE2), Italy (IT4), and Norway (NOR2).**(A)** Principal component analysis of CGN, CHG, and CHH methylation in ecotypes grown at 18 ºC (blue symbols) and 28 ºC (red symbols). Red lines indicate the x, y and z axes. **(B)** Methylation differences between temperatures (28 vs. 18 ºC) on *F. vesca* chromosome 3 (Fvb3) in the different ecotypes. Methylation differences were calculated for 50 kb windows using 18 ºC as a reference. **(C)** Methylation patterns along all seven *F. vesca* chromosomes for all ecotypes and both temperatures, using a 50 kb window. Lanes 1-8 show methylation levels for each ecotype-temperature combination. Connecting lines in the middle indicate coding regions (CDS) regions that showed synteny.

On a chromosome scale, all ecotypes had similar methylation patterns for the CGN context at a 50 kb-window resolution, whereas there were clear differences between ecotypes for the CHG and CHH contexts at this resolution (Fig. 3B, Fig. S10). Compared to the other ecotypes, NOR2 had a higher methylation increase at 28 vs. 18 ºC for all methylation contexts and all chromosomes, with an average of ∼4%, ∼5%, and ∼1% hypermethylation in the CGN, CHG, and CHH context, respectively (Table S5). A detailed analysis of all possible trinucleotide methylation contexts suggested that temperature treatment introduced no obvious methylation bias of the nucleotide contexts (Fig S11, S12, S13). This aspect was therefore not included in further analyses.

We compared methylation responses to temperature treatment between all ecotypes by plotting heat-maps of chromosomes, ecotypes, and temperature treatments for all three methylation contexts. In all ecotypes, significant changes in hypo- and hypermethylation occurred along the chromosomes, and the largest treatment-specific methylation increases at 28 ºC were observed for the CHH context (Fig. 3C). For the CGN and CHG contexts the most striking hypo- and hypermethylated areas were largely shared between ecotypes and temperature treatments, although there was some treatment-specific hypermethylation for the CHG context in all ecotypes (e.g. in the middle of chromosomes Fvb5 and Fvb6 at 28 ºC; Fig 3C). In rare cases, we observed ecotype-specific treatment effects, for instance, at the end of chromosome Fvb1 and in the middle of Fvb6 in the CHG context in NOR2 and at the beginning of Fvb7 in the CHH context in IT4 (Fig. 3C).

### Methylation of genomic features differs between temperature treatments

To determine the association between DNA methylation and genes, REs, and pseudogenes we calculated DNA methylation patterns along these genomic features, as well as 2 kb upstream and downstream of them. For all genic methylation contexts, methylation levels were lowest around transcription start sites (TSS) and transcription termination sites (TTS), as previously reported (Fig 4A, Fig S14) (Edger *et al.* 2018; Liu *et al.* 2022). For REs, we observed a plateau of hypermethylation in the RE body region, flanked by regions with decreasing methylation levels for all contexts (Fig 4B, Fig S14). For pseudogenes, methylation levels peaked around start and end sites and were lower within body regions (Fig 4C, Fig S14). Methylation levels were higher in plants propagated at 28 vs. 18 ºC in all genomic features, both in body regions and flanking regions. Thus, on average, DNA methylation increased with increasing temperature (Fig 4 A,B, Fig S14).

**Fig 4.**
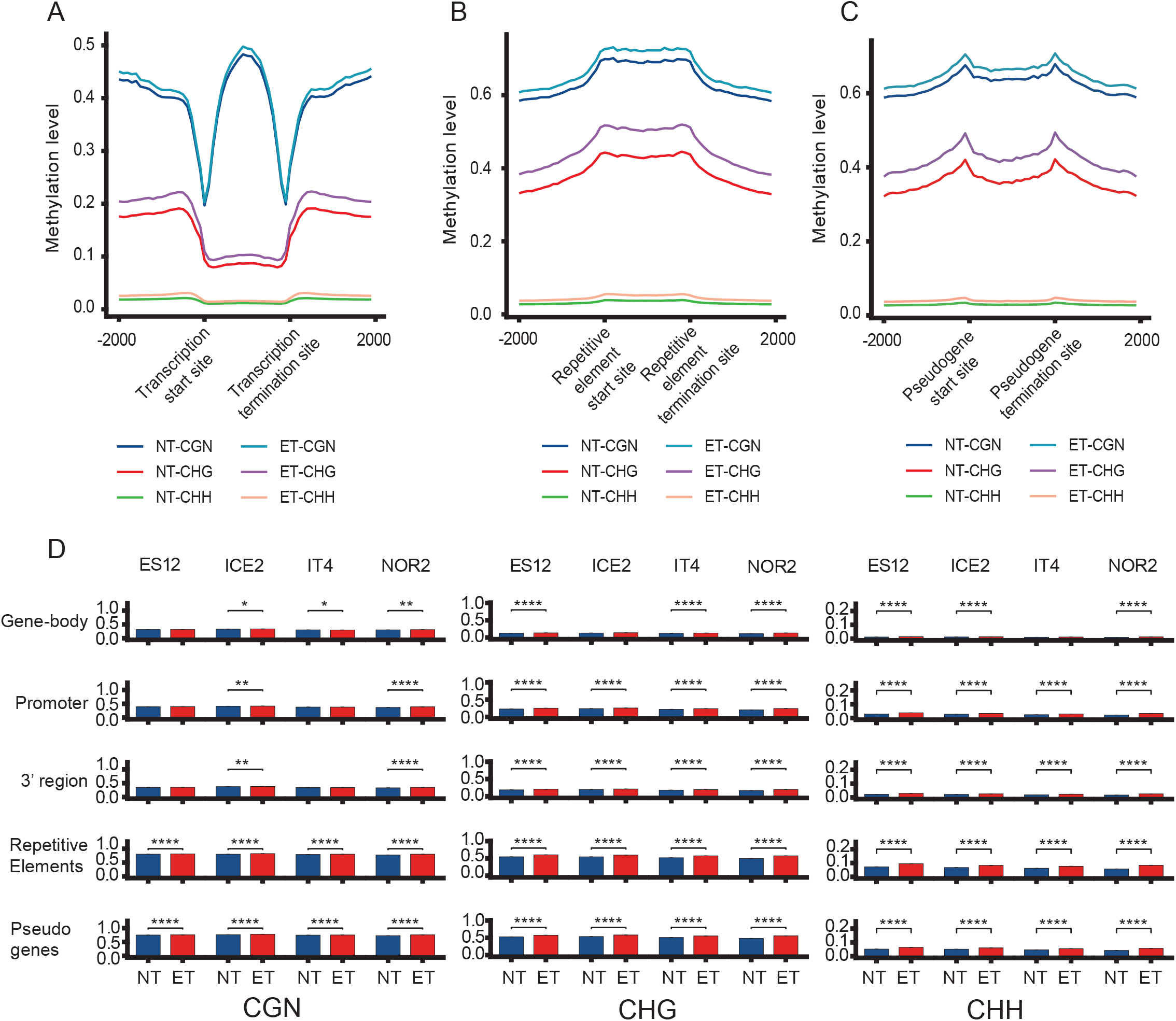
DNA methylation landscape of protein-coding genes, repetitive elements, and pseudogenes in leaves of different *Fragaria vesca* ecotypes propagated at normal (18 ºC) and elevated (28 ºC) temperature conditions. Methylation pattern of protein coding genes**(A)**, repetitive elements **(B)**, and pseudogenes **(C)** in the NOR2 ecotype. Each genomic feature and regions 2 kb upstream and downstream to these were divided into 20 pieces to calculate methylation levels (methylated reads/total reads). Colored lines show different combinations of temperature and methylation contexts. **(D)** Average methylation level of different genomic features in four *F. vesca* ecotypes (ES12, ICE2, NOR2 and IT4) for different methylation contexts (CGN, CHG, and CHH) and temperature conditions (18 and 28 ºC). Asterisks indicate significant differences: * 0.01 ≤ p < 0,05; ** 0.001 ≤ p < 0.01; *** 0.0001 ≤ p < 0.001; **** 0.00001 ≤ p < 0.0001. Error bars show 95% confidence intervals.

We compared global methylation levels in genes, REs, and pseudogenes to understand the hypermethylation observed in plants propagated at 28 ºC. On average, methylation levels in the CGN context were approximately 30%, 40%, and 35% for promoter, gene-body and 3’-primed regions, respectively. CGN methylation levels were considerably higher in REs and pseudogenes (80% and 75%, respectively) (Table S6). REs and pseudogene classes were significantly hypermethylated in all methylation contexts and all ecotypes (Fig 4D, Table S6). Notably, the NOR2 ecotype also showed significant hypermethylation in genes. Plants propagated at 28 ºC also showed significant hypermethylation in the CHG and CHH contexts in most ecotypes. Exceptions were gene body methylation in the CHG context in ICE2 and in the CHH context in IT4. For the CGN context, ecotypes differed more, although both NOR2 and ICE2 showed significant hypermethylation (Fig 4D, Table S6).

### Regions with differential CHG and CHH methylation are generally hypermethylated

We identified differentially methylated regions (DMRs) between ecotypes that were propagated at 18 vs. 28 ºC for three asexual generations (AS3) (Fig 5A, Fig S15, Table S7). As a control we identified DMRs between plants propagated at 18 ºC for three vs. one asexual generation (AS3 vs. AS1), to exclude the effect of temperature (Fig S16, S17, Table S8). Plants showed minor changes in DNA methylation across two asexual generations at 18 ºC. The total number of DMRs observed in the trans-generational control (AS3 vs. AS1) was significantly lower than that in the temperature comparison (18 vs. 28 ºC; 0.05 > p > 0.001) (Table S9). Separate analyses of hyper- and hypomethylated DMRs showed that hypermethylated DMRs in the CHG and CHH contexts were significantly more common in the temperature comparison than in the trans-generational control (p < 0.008 and p < 0.001 for CHG and CHH, respectively). Hypomethylated DMRs in the CGN context were also significantly overrepresented in the temperature comparison (p < 0.018) (Table S9). This suggested that temperature treatment had a significantly larger impact on the methylome than changes accumulated across two asexual generations.

**Fig 5.**
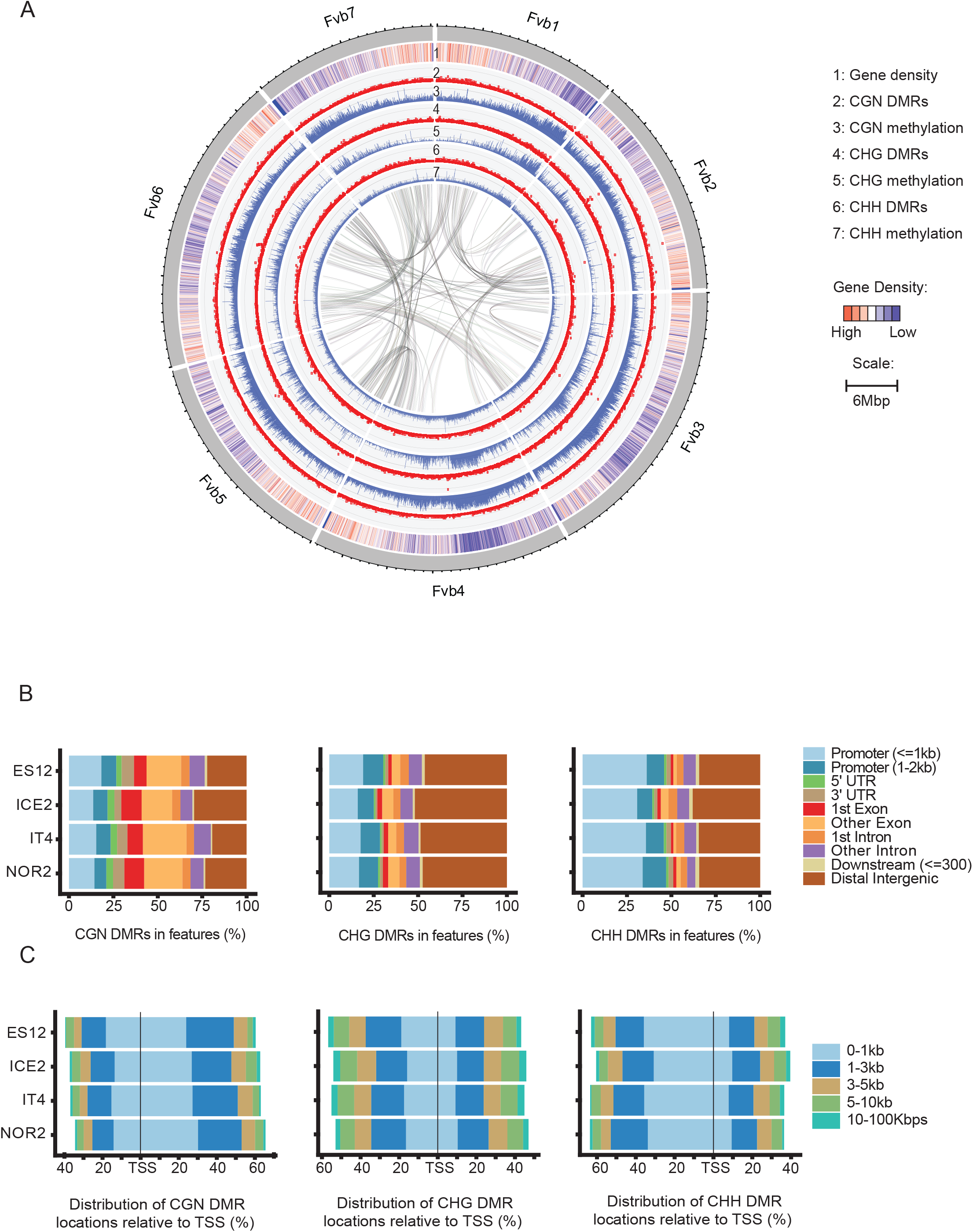
Chromosomal and genic distribution pattern of differentially methylated regions (DMRs) in *Fragaria vesca* leaves. **(A)** Pattern of DMRs on each *F. vesca* (Fv) chromosome in the NOR2 ecotype. Lanes 1-7 show gene density, DMR density and methylation level (%) for different methylation contexts (CGN, CHG, and CHH). Connecting lines in the middle indicate coding regions (CDS) regions that showed synteny. **(B)** Distribution of DMRs for different methylation contexts across different genomic features in four F. vesca ecotypes. **(C)** Distribution of DMRs for different methylation contexts according to their distance (5’-3’ direction) from transcription start sites (TSS) in different ecotypes.

The number of DMRs for different methylation contexts varied between ecotypes. In the temperature comparison (18 vs. 28 ºC), we identified a total of 738-1538 CGN DMRs, 2609-4799 CHG DMRs, and 625-1023 CHH DMRs in the four ecotypes (Table S9). NOR2 had the greatest number of DMRs for all three methylation contexts and CHG methylations were most common in all ecotypes (Table S9). Furthermore, most DMRs in the CHG and CHH contexts were hypermethylated at 28 ºC. In contrast with the CGN context, which typically was hypomethylated, more than 98% of all CHG DMRs and 94% of all CHH DMRs were hypermethylated at 28 ºC, demonstrating that hypermethylation was the main methylation response to increasing temperatures and that hypermethylation occurred both in symmetric (CHG) and asymmetric (CHH) contexts.

### Differentially methylated regions pinpoint temperature treatment-specific methylation peaks

By plotting DMRs for different methylation contexts across all chromosomes we found that DMRs had a distinct distribution pattern on each chromosome, with several regions with a high density of DMRs. Some DMR regions were unique to certain ecotypes and others were shared between ecotypes (Fig 5A, Fig S17, Table S10).

Most CGN and CHH DMRs were located in gene body and promoter areas, whereas most CHG DMRs were found in intergenic regions (Fig. 5B). For the CGN context, the proportion of DMRs was greater in gene bodies than in promoter areas, while the opposite pattern was found for the CHH context (Fig 5B). About 50% of CHG DMRs were localized in intergenic areas in NOR2, whereas the corresponding fraction in ES12 was ∼25% (Fig. 5B). While most CHG and CHH DMRs were situated in the 3’ direction relative to the transcription start site, CGN DMRs tended to be located in the 5’ direction (Fig. 5C).

We performed a HOMER2 analysis to investigate if DMRs correlated with known sequence motifs (Heinz *et al.* 2010). Interestingly, motif identification in 200 bp regions adjacent to CHG DMRs revealed an overrepresentation of the MYB motif (Table S11, S12).

Using a 50 kb window, we identified eight DMR peak regions in the CHG context that were shared in all ecotypes. These regions were located on Fvb2, Fvb3, Fvb5, and Fvb6 (Table S10) and were adjacent to, but did not completely overlap with, methylation peaks on each chromosome. Overall, the DMRs in each peak region only occupied a small region of each 50 kb window.

To study the identified DMR peak regions in more detail we plotted the chromosomal locations of CHG DMRs against different genomic features. The number of genes identified within regions designated as DMR peak regions varied between two and eight (Table S10), but the majority of DMRs in the DMR peak regions did not overlap with gene-body regions and did not show a strong correlation with genes or other genomic features located in DMR peak regions (Fig 6). Except for a single region in Peak 4, all DMRs in peak regions were hypermethylated (Fig 6). Strikingly, the differential methylation patterns in both hypo- and hypermethylated DMR peaks were conserved between ecotypes, suggesting a common underlying methylation mechanism. A Gene Ontology (GO) analysis of genes located in DMR peaks identified several GO terms related to the regulation of microtubule cytoskeleton organization that were enriched >250-fold (p_adj_ < 0.000002, Table S10). All the identified genes in this category were *TARGET OF PROTEIN FOR XKLP2 (TPX2)* genes. A closer inspection revealed a total of five *TPX2* genes (out of 21 in the whole genome) located in three DMR peaks (Peak 3, 6, 7; Fig 6, Table S10). Thus, *TPX2* genes were significantly enriched in DMR peaks that were shared between ecotypes.

**Fig 6.**
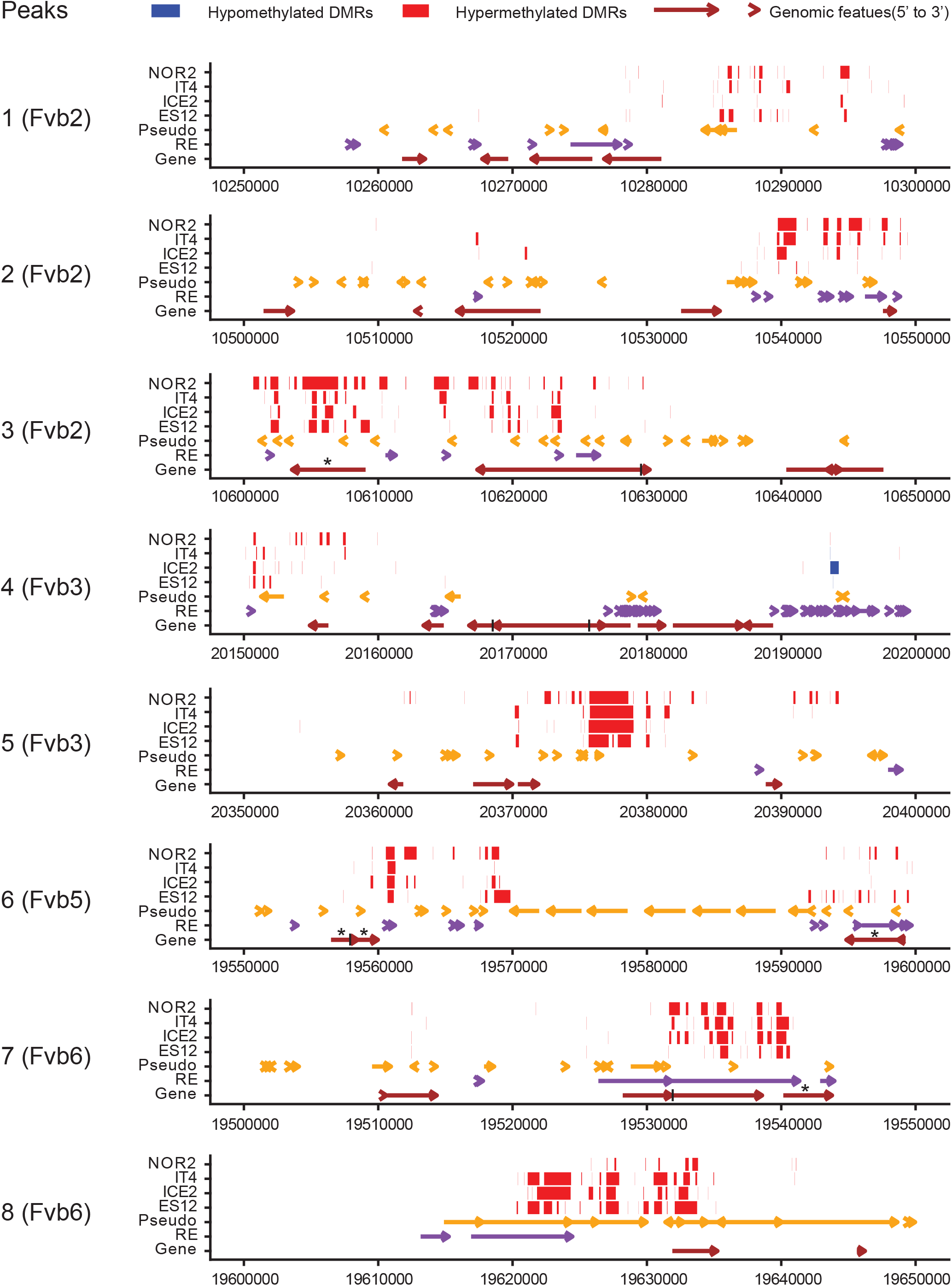
Differentially methylated regions (DMRs) and genomic features inside DMR peaks in leaves of different *Fragaria vesca* ecotypes. Distribution of DMRs, pseudogenes, repetitive elements (REs), and genes along eight 50 kb wide DMR peaks for the CHG methylation context in four *F. vesca* (Fv) chromosomes. The x-axis indicates location along the DMR peak. Red and blue bars represent hyper- and hypo-methylated DMRs, respectively. Coloured arrows indicate pseudogenes (yellow), REs (purple), and genes (brown). Asterisks indicate predicted TPX genes.

### Differentially expressed and differentially methylated genes (DEDMGs) are detected in all four ecotypes

We defined differentially methylated genes (DMGs) as genes that had DMRs located in their promoter and/or gene body (Table S13). Most DMGs were methylated in the CHG context, whereas the CGN and CHH contexts were less prevalent. Each ecotype had its own unique set (>1144) of DMGs, and only a small number of DMGs with the same methylation context was shared between all ecotypes (15 in CGN, 85 in CHG, and 14 in CHH context) (Fig 7A, Table S13). If we define DMGs as being differentially methylated in any context, the number of shared DMGs across all ecotypes was slightly higher (167, Figure 7A, Table S13).

**Fig 7.**
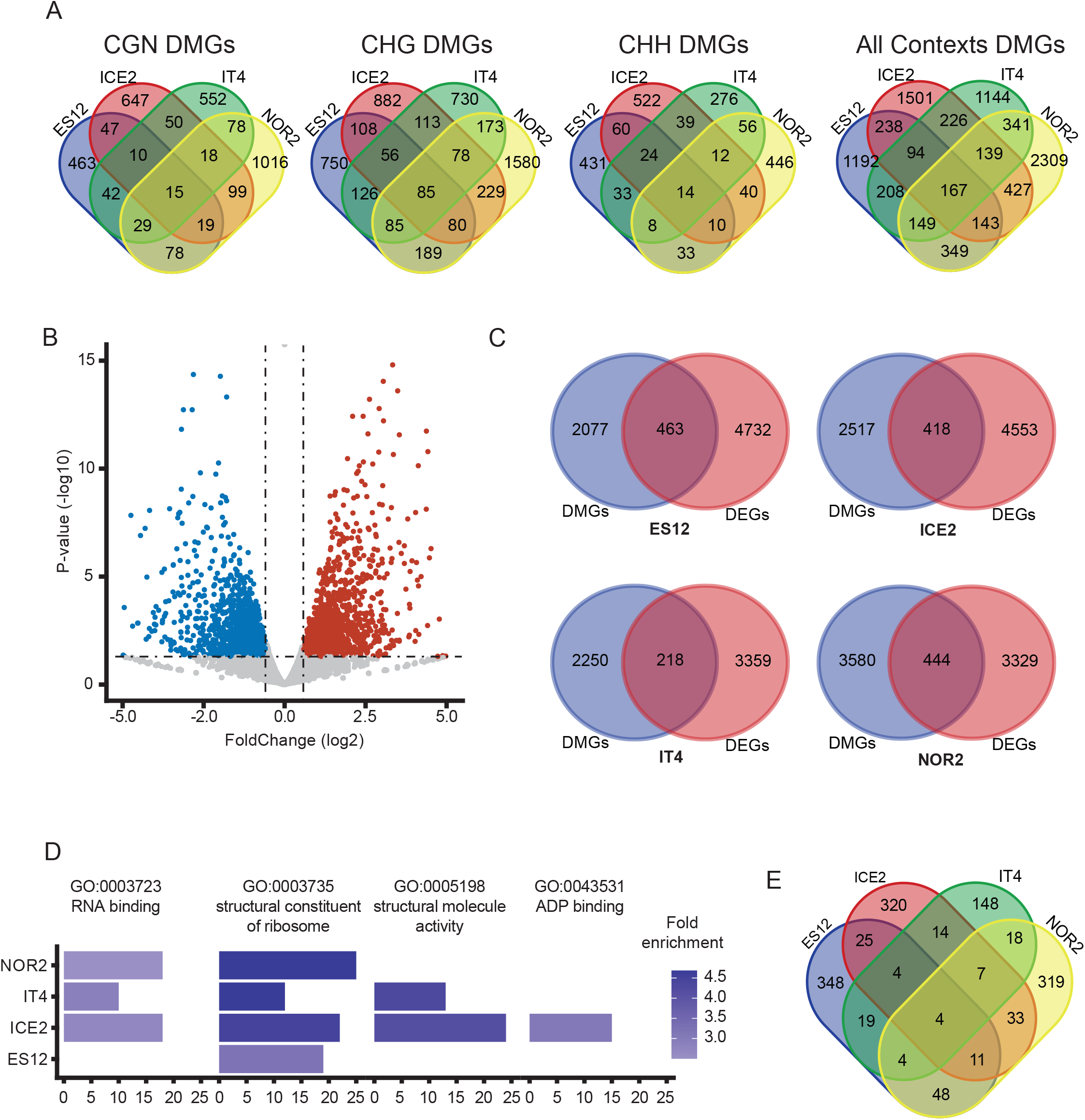
Effects of DNA methylation on gene expression in *Fragaria vesca* ecotypes propagated under different temperature conditions. **(A)** Venn diagrams showing differentially methylated genes (DMGs) for different methylation contexts (CGN, CHG, CHH) and ecotypes (ES12, ICE2, NOR2, IT4). **(B)** Volcano plot showing differentially expressed genes (DEGs) in NOR2 plants propagated at 28 ºC vs. 18 ºC. Blue and red dots show down- and upregulated genes, respectively. Dashed lines delineate P-value ≤ 0.05 and log_2_FoldChange ≥ 1.5. **(c)** Venn diagrams showing overlap between DEGs and DMGs in the different ecotypes. **(D)** Enriched Gene Ontology (GO) terms among differentially expressed differentially methylated genes (DEDMGs) in the different ecotypes. The x-axis indicates the number of DEDMGs per GO term. Color shading indicates fold enrichment at 28 vs. 18 ºC. (E) Venn diagrams showing DEDMGs in all ecotypes.

We quantified changes in the expression of DMGs by first identifying differentially expressed genes (DEGs) in all ecotypes (Fig 7B, Fig S18) and then comparing DMGs to DEGs. Compared to a random prevalence, we found a significant over-representation of differentially expressed DMGs (DEDMGs) in plants propagated at 28 ºC (10-20% DEDMGs, Fisher exact test, Fig 7C, Table S14). Across all ecotypes, DEDMGs were enriched for four GO terms: RNA binding, structural component of the ribosome, structural molecular activity, and ADP binding (Fig 7D, Table S15). Among these, ‘structural component of the ribosome’ was shared by all ecotypes and ‘RNA binding’ was shared by all except ES12 (Fig 7D, Table S15).

Individual ecotypes had 218-463 DEDMGs and 68-76% of these were ecotype-specific (Fig 7C). Nonetheless, some DEDMGs were shared between ecotypes and four DEDMGs were shared by all ecotypes (Fig 7E). These shared DEDMGs encoded a SERPIN family protein serine protease inhibitor, a tubulin alpha-5 protein, a DNA-binding protein HEXBP-like, and a sieve element occlusion amino-terminus protein.

To characterize these four shared DEDMGs in more detail we plotted their gene body and promoter regions together with the location of DMRs and compared these data with gene expression. In all shared DEDMGs, the expression change was always going in the same direction in all four ecotypes. Three of the shared DEDMGs were hypermethylated and upregulated, whereas one was hypomethylated and downregulated at 28 vs. 18 ºC (Fig 8). Hypermethylated and upregulated DEDMGs were methylated in the CHG and CHH contexts in regulatory regions, whereas the hypomethylated and downregulated DEDMG was methylated in the CGN context in the gene body (Fig 8). Strikingly, the differential methylation patterns were very similarly positioned across ecotypes in all hypo- and hypermethylated DEDMGs, except for the SERPIN family protein FvH4_5g10280.

**Fig 8.**
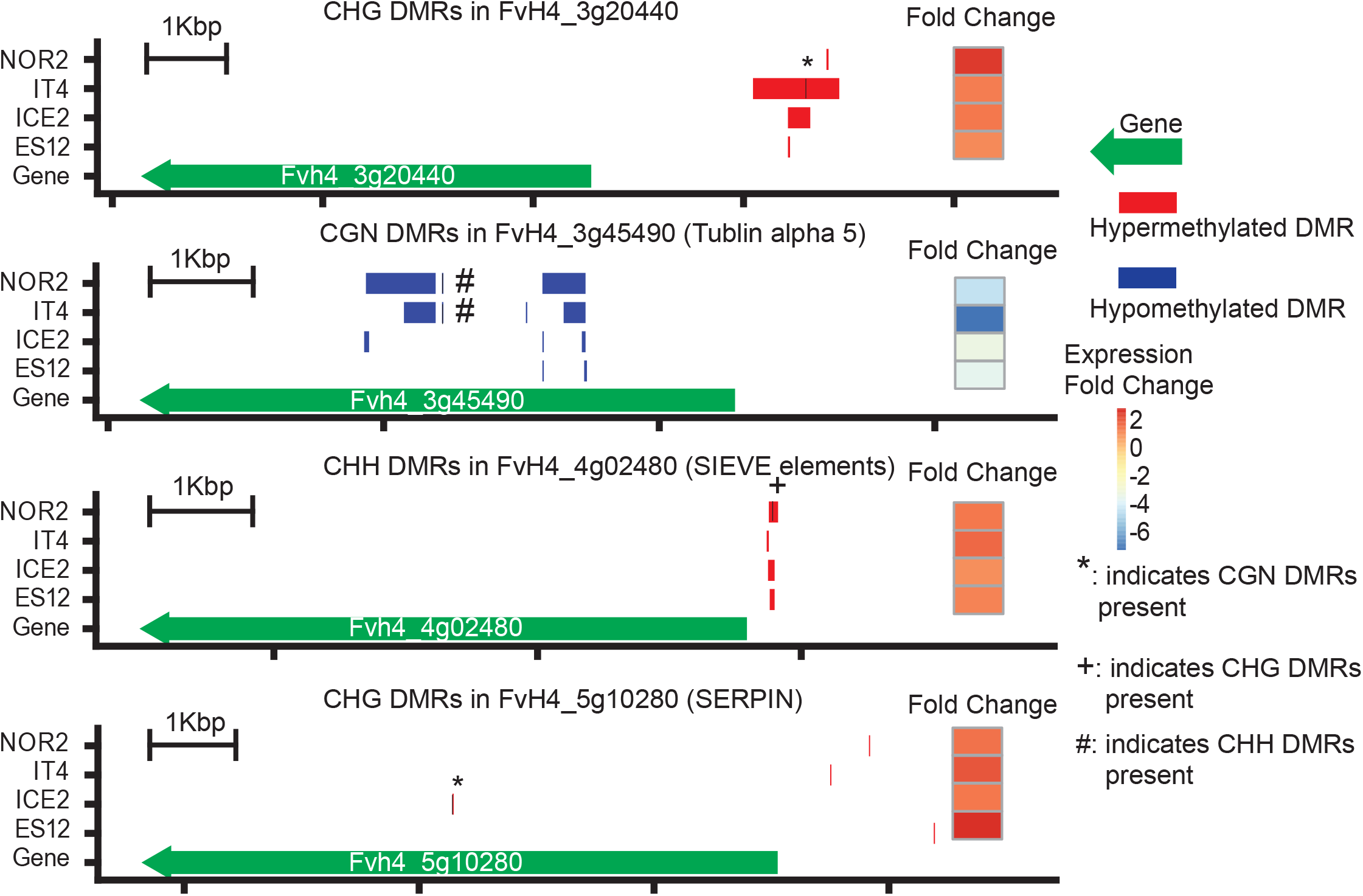
Functional analysis of differentially expressed and differentially methylated genes (DEDMGs) in *Fragaria vesca* ecotypes propagated at normal (18 ºC) and elevated (28 ºC) temperatures. Expression and methylation status of four DEDMGs on *F. vesca* chromosome 4 (FvH4) that are shared by all four ecotypes (ES12, ICE2, NOR2, IT4). Red and blue bars represent hyper- and hypomethylated differentially methylated regions (DMRs), respectively, in different methylation contexts (CHG, CGN, CHH). *, +, and # indicate presence of CGN, CHG, and CHH DMRs, respectively, at the position indicated by the black vertical line.

### Ecotypes show unique methylation and gene expression patterns in specific functions and pathways

Having described DEDMGs shared by all ecotypes we went on to characterize ecotype-specific changes in the epigenetic machinery as well as in flowering-, temperature- and gibberellin-related genes and transcription factors. For these specific pathways we found a total of 1318 DEDMGs with an absolute fold-change > 1.5 (p_adj_ < 0.05) in three or fewer ecotypes (Table S14). Increasing the stringency to an absolute fold-change >2.0 limited the number of DEDMGs to 678 (Table S14).

Methylation and expression of several genes in the epigenetic machinery changed in plants propagated at 28 ºC (Table S14, Fig S19). The ES12 and IT4 ecotypes shared upregulation of a gene encoding a chromatin remodeling complex SWIB/MDM2, Plus-3, and GYF domain-containing protein (FvH4_1g05690), whereas ES12 and NOR2 shared regulation of a chromatin reader Tudor-domain SAWADEE homolog (FvH4_6g40530) involved in RNA-directed DNA methylation, as well as a small RNA-degrading nuclease (Fig S19). Other differentially regulated genes in individual ecotypes included a gene in the structural maintenance of chromosomes (SMC) family (FvH4_7g27644) that participates in higher-order chromosome organization and dynamics (in ES12), a homolog of the XH/XS domain-containing protein (FvH4_3g36253) involved in dsRNA binding in RdDM-related siRNA biogenesis (in ICE2), another RdDM-related SAWADEE protein homolog (FvH4_3g20410 in IT4), and a SETD group SET domain-containing protein gene and a PHD finger protein gene (FvH4_7g20720 and FvH4_4g12240 in NOR2) (Fig S19).

Because temperature affected flowering time in NOR2 and ICE2 ecotypes, we examined changes in methylation and expression of genes homologous to Arabidopsis flowering time genes (according to the FlorID database) and other genes annotated as putative flowering-related genes (Fig S20). The ICE2 and NOR2 ecotypes shared regulation of a HY5 homolog (FvH4_4g21800) that, based on findings in Arabidopsis, is putatively involved in the epigenetic regulation of flowering (Chu, Yang, Zhuang, Gao & Luo 2022). These ecotypes also shared the downregulation of a TCP transcription factor-homolog, a putative interactor of FLOWERING LOCUS T (FT, FvH4_6g00090) (Ho & Weigel 2014). Interestingly, two *JOINTLESS-like/SVP/AGL22* homologs (FvH4_5g35400 and FvH4_5g35401) were downregulated in ES12, while a related AGL24 homolog (FvH4_3g03630) was upregulated in ICE2. Both genes are involved in SOC1-dependent regulation of flowering in Arabidopsis (Lee & Lee 2010). Four homologs of homeobox-containing protein genes linked to flowering were also identified, two in ES12 (FvH4_3g16431 and FvH4_5g27650) and two in ICE2 (FvH4_4g09960 and FvH4_7g23561). In ES12, we also found a gene encoding a circadian clock-related B-box-containing protein (FvH4_6g43580), an *VERNALIZATION* homolog (VRN1, FvH4_7g07610), and a *WRKY70-like* homolog (FvH4_7g26030) (Table S14).

We also detected a total of 13 temperature-related DEDMGs, including five encoding chaperones, two with protease activity, and two involved in MEDIATOR of RNA polymerase II subunit. Six of the 13 DEDMGs were shared by pairs of ecotypes (ES12-NOR2 and ICE2-NOR2). However, the remaining DEDMGs were found only in one ecotype (Table S14, Fig S21), suggesting that these temperature-related DEDMGs do not represent a general response to the temperature treatment.

Since the gibberellin pathway contributes to both flowering and stolon formation (Andrés *et al.* 2021), we examined the differential expression and methylation patterns of gibberellin genes. Interestingly, ICE2, IT4, and NOR2 shared a DEDMG encoding a CYP72A15 homolog (FvH4_3g17260) that catalyzes the 13-hydrolyzation of gibberellic acid (GA) and reduces GA activity (He *et al.* 2019). NOR2 also contained an additional DEDMG CYP72A15 homolog (FvH4_7g17900). Other GA-related DEDMGs were unique to specific ecotypes: a GA receptor (FvH4_2g32670), a GA-regulated protein (FvH4_2g24370), and a GA 2-oxidase (FvH4_3g05530) DEDMG in ICE2, a GA-regulated protein (FvH4_5g01680) in IT4, and a GA-STIMULATED ARABIDOPSIS 6 homolog (FvH4_5g14950) in NOR2 (Table S14).

### Repetitive elements impact the expression of nearby genes

Repetitive elements (REs) are well known targets of DNA methylation and RE methylation may affect the expression of nearby genes. We therefore examined the methylation status of REs in the *F. vesca* genome, starting by comparing DMRs across temperature treatments. Most DMRs within REs were hypermethylated in plants propagated at 28 vs. 18 ºC, particularly in the CHG and CHH contexts (Fig 9A). Hypomethylation was very rare in these methylation contexts. In the CGN context, NOR2 had the highest degree of hypermethylation among the four ecotypes whereas ES12 had more hypomethylation than other ecotypes. Hypermethylation in NOR2 was paramount in all methylation contexts, indicating increased silencing of REs in response to elevated temperatures in this ecotype (Fig 9A).

**Fig 9.**
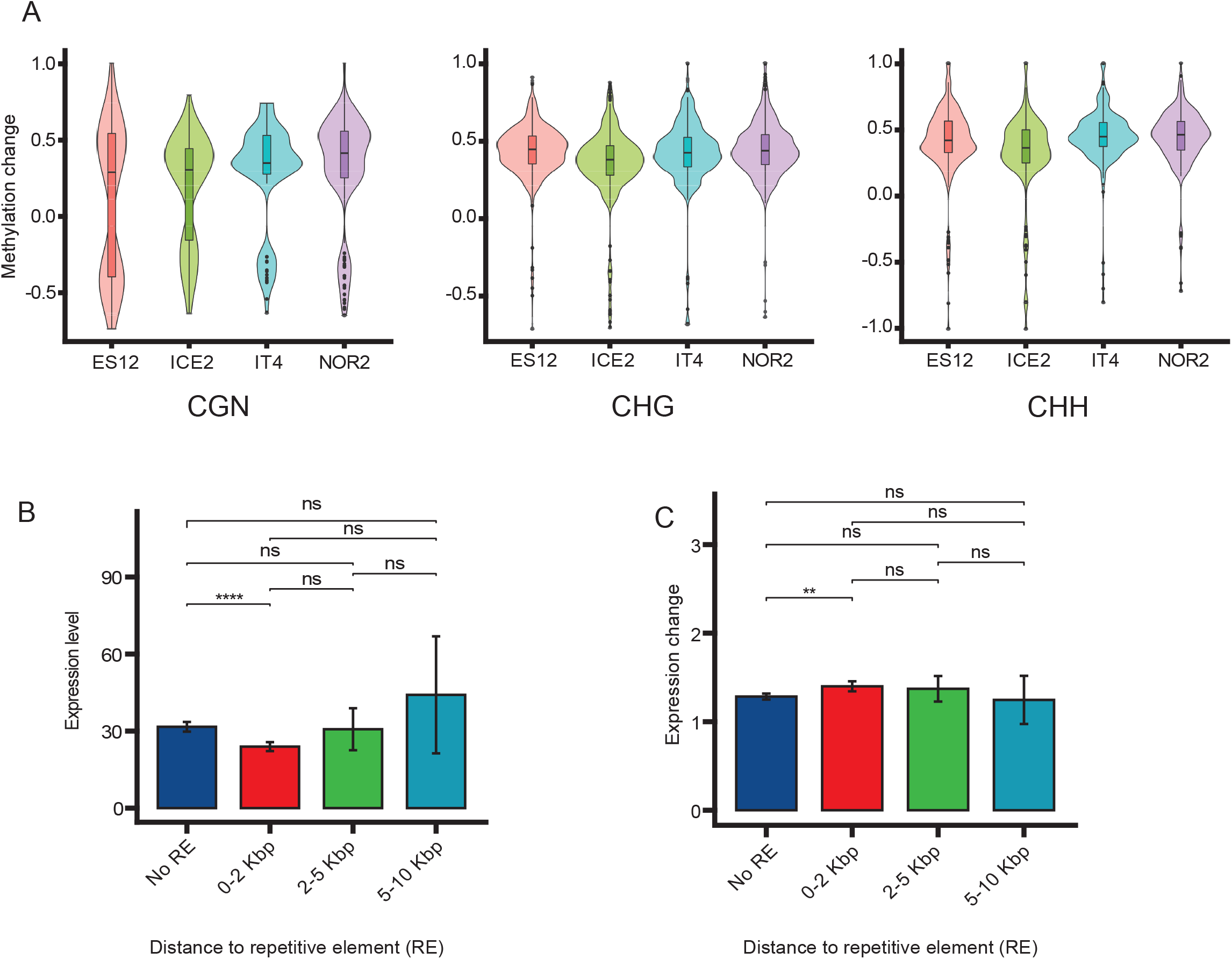
The impact of methylation of repetitive elements (REs) on gene expression in *Fragaria vesca* ecotypes propagated at normal (18 ºC) and elevated (28 ºC) temperatures. **(A)** Violin plots showing methylation change in REs in different ecotypes (ES12, ICE2, NOR2, IT4) for the CGN, CHG, and CHH methylation contexts. The average methylation value in differentially methylated regions was calculated based on the methylation level of each methylated site, discarding any unmethylated sites. The methylation value was calculated as methylated reads/total reads. Violin plots show the median (horizontal black line), 25 and 75 quantiles (colored bars), outliers (black circles), and 95% credential intervals (error bars). **(B-C)** Positional effect of REs on gene expression level **(B)** and absolute gene expression change **(C)** in ecotype NOR2. The distance from the gene to the nearest RE is given on the x-axis. Asterisks indicate significant difference calculated by games_howell_test: ** 0.001 ≤ p < 0.01; **** 0.00001 ≤ p < 0.0001. Error bars show 95% confidence intervals.

To determine if REs affected the expression of nearby genes, we classified genes into four categories based on the distance between the gene and the closest RE: (1) no REs within 10 Kbp, (2) an RE within 2 Kbp, (3) an RE within 2 to 5 Kbp, and (4) an RE within 5 to 10 Kbp. Only REs within 2 Kbp from genes had a significant positional effect on transcription (Games-Howell test, 0.05 > p > 0.001) (Fig 9B, Fig S19A). The general effect of having an RE 2 Kbp or less from a gene was a significantly reduced transcript level of that gene (Fig 9B, Fig S19A). Genes with REs close-by had a significantly greater change in transcript level than genes with no REs nearby (DEG at 28 vs. 18 ºC, abs_log2FC, Games-Howell test, 0.001 ≥ p-value ≥ 0.005) (Fig 9C, Table S16, Fig S19B). There was a significant location effect of REs in all ecotypes, causing transcriptional changes in response to elevated temperatures that likely were due to increased DNA methylation of REs.

## Discussion

This is the first study of how conditions experienced during asexual reproduction influences the epigenetic memory in *F. vesca*. By exploring methylomes and transcriptomes from young leaves of four ecotypes we describe DNA methylation changes and their putative effects on gene expression, and explore correlations to phenotypic changes. The *F. vesca* ecotypes we studied originate from very different habitats in Spain, Italy, Iceland, and northern Norway and exhibit distinct phenologies. Using these different ecotypes, we analysed methylome signatures correlated to phenological changes induced by exposing plants to different temperature conditions during asexual propagation. Our data can help us further understand the evolutionary adaptations of perennial plants to a rapidly warming world, and how the methylome and transcriptome contribute to phenological plasticity.

### Temperature treatment induces stable plant phenotypes

The phenology of *F. vesca* was affected by temperature during asexual reproduction. Elevated temperatures (28 ºC) during the vegetative phase delayed the time to flowering in the high-latitude ecotypes NOR2 and ICE2, and the impact increased from the first to the third asexual generation. Epigenetic memory to temperature conditions experienced during asexual reproduction have been observed previously in gymnosperm plants. Significantly different temperature sums experienced during Norway spruce embryogenesis (a form of asexual reproduction) induce epitypes with lasting changes in adult tree traits such as bud phenology and frost tolerance in a reproducible and predictable manner (Kvaalen & Johnsen 2008; Carneros *et al.* 2017). Although the effect of temperature on flower induction of *F. vesca* is well established (Rantanen *et al.* 2015), this report is the first indication of an epigenetic memory-response caused by temperature during asexual reproduction. The low-latitude ecotypes IT4 and ES12 did not flower in the first asexual generation and did not respond significantly to temperature treatment. The shorter time to flowering and higher plasticity in NOR2 and ICE2 may be an adaptation to shorter growth periods at high latitude. The fact that the altered phenotype was detectable after three asexual generations indicates that effects on flowering time may accumulate over several generations with repeated treatment.

The impact of temperature on stolon formation (in ES12 and IT4) and petiole length (in ES12, ICE2 and NOR2) was strongest in the first asexual generation but remained significant for three generations in IT4 (for stolon number) and NOR2 (for petiole length). Thus, there was a possible epigenetic impact of temperature treatment on these adaptive features but it was weaker than for flowering. Taken together, variable responses to treatment in different ecotypes suggest that ecotypes differ in phenotypic plasticity. NOR2 was the most plastic ecotype with significant responses in both flowering time and petiole length.

### Temperature treatment causes massive changes in DNA methylation

We used whole genome bisulfite sequencing to probe changes in the methylomes of the four ecotypes following temperature treatments. Individual hyper- and hypomethylated peaks occurred in all ecotypes, indicating that there were numerous global and specific methylation differences between ecotype methylomes. When comparing the temperature-induced methylation changes for all genes and repetitive elements, NOR2 showed the largest change. This suggests that NOR2 has the most plastic methylome of the four ecotypes and mirrors the higher phenotypic plasticity observed in NOR2. Indeed, NOR2 also showed significant methylation changes in all genic features (gene body, promoter, 3’ region) in all methylation contexts. In repetitive elements (RE) and pseudogenes, a significant response to temperature was found in all four ecotypes for all three methylation contexts, which contrasts with the more subtle and selective methylation changes observed in other genic features. A similar response has been described in mulberry (*Morus notabilis*) and *Arabidopsis* following pathogen infection (Dowen *et al.* 2012; Xin *et al.* 2021), suggesting a common methylation response to both biotic and abiotic impacts.

We found that CHG and CHH DMRs mostly were located in intergenic regions and in the 5’ direction of the transcription start site (TSS), whereas CGN DMRs mostly were located in promoter and gene-body regions. In plants, the gene body region is usually enriched with CGN methylation whereas CGN methylation is depleted at the transcriptional start and termination sites. Similar to our findings, genes with high CGN gene body-methylation are usually highly expressed and conserved (Tran *et al.* 2005; Takuno & Gaut 2012, 2013; Seymour, Koenig, Hagmann, Becker & Weigel 2014; Bewick *et al.* 2016; Bewick & Schmitz 2017)).

Regardless of ecotype, the CHG methylation context had most DMRs and DMGs in our study. In contrast to CGN methylation, which showed both hypo- and hypermethylation, most treatment-induced CHG and CHH DMR’s were hypermethylated in all ecotypes. This indicates that more than one methylation pathway was triggered by the temperature treatment. In plants, CHG and CHH methylation and maintenance are mediated by the DNA methyltransferases CMT3 (CHG) and CMT2 and DRM1/DRM2 through RdDM (CHH) (Erdmann & Picard 2020), indicating that both pathways are involved in plant responses to temperature treatment.

We speculate that induced hypermethylation in the CHG and CHH contexts could be due to “self reinforcing” chromatin interactions. In one potential scenario the histone methyltransferases KRYPTONITE (KYP)/(SU(VAR)3-9) HOMOLOGUE 4 (SUVH4) and its homologs SUVH5/6 could recognize methylated CHG or CHH and lead to targeted methylation of H3K9me2 (Jackson *et al.* 2002, 2004; Malagnac *et al.* 2002; Johnson *et al.* 2007). CMT2/3 is recruited to H3K9me2 and methylates nearby CHG or CHH sites, whereas methylated CHG and CHH in turn recruit KYP and SUVH5/6 leading to reinforcement of both histone H3K9me2 and CHG/CHH DNA methylation. Furthermore, RELATIVE OF EARLY FLOWERING 6 (REF6) and putatively also other Jumonji domain-containing (Jmjc) histone demethylases, are repressed by non-CGN methylation (Miura *et al.* 2009; Li *et al.* 2016; Qiu *et al.* 2019), further reinforcing histone H3K9me2 and enhancing CHG/CHH DMR rich chromosomal regions.

### DMR peaks shared between ecotypes identify the target of XLP2 genes

We observed several regions with a high density of DMRs, denoted DMR peaks (DMRP). Most DMR-rich regions differed between ecotypes, but shared DMRP existed on chromosomes Fvb2, Fvb3, Fvb5, and Fvb6. Zooming in on these DMRP, we found only eight that were shared by all four ecotypes and these DMPR were exclusively in the CHG context. We found five *TARGET OF XKLP2* (TPX2) genes located in three peaks, and three of these TPX2s have homologs in *Arabidopsis*. TPX2 is a key mitotic spindle-assembly factor and acts like a spindle regulator. The *Arabidopsis* TPX2 homolog WAVE-DAMPENED2, has been found to reduce the waviness of root growth (Smertenko *et al.* 2021). Although TPX2 is involved in spindle formation, its homologs have evolved into many roles. They function in hormonal regulation of hypocotyl cell elongation, light signaling, vascular development, and abiotic stress tolerance (Smertenko *et al.* 2021). Whether increased methylation of TPX2 genes is a direct or indirect effect in that cell division was impacted by temperature treatment, remains speculative since TXP2 expression was only impacted in three out of four ecotypes. It is, however, interesting that five (out of 21) members of the same protein family were situated within DMR peaks that make up less than 0.2% of the *F. vesca* genome (0.4Mb/240Mb). The high methylation level may suggest that these regions, displaying normal CG levels, contain common recognition features that are targeted by the epigenetic machinery following temperature treatment. The observation that all DMRP were enriched in the CHG methylation context may propose DMRP as a future model system to uncover mechanistic pathways by which environmental stressors target CHG DNA methylation.

### Methylation responses to temperature treatments are accompanied by transcriptional change

Identification of DMRs allowed us to call ∼2500 to ∼4000 treatment-induced differentially methylated genes (DMGs) in the different ecotypes, most of which were associated with increased CHG methylation. We noted that the identified DMGs were predominantly ecotype-specific, in line with the observation that the phenotypic responses we observed also were ecotype-specific. The NOR2 ecotype had the highest number of DMGs, correlating with the observation that NOR2 displayed the strongest induced methylation response to treatment.

Temperature treatment caused transcriptomic changes ranging from ∼3500 to 5000 differentially expressed genes (DEGs) in the different ecotypes. We found that 10 to 20% of these DEGs were also DMGs, denoted as DEDMGs. These DEDMGs were statistically overrepresented compared to random gene samples, suggesting that induced DNA methylation impacted the expression of at least a subset of these DEGs. The finding that translation-related GO terms were enriched in DEDMGs in all four ecotypes hints that the process of translation was impacted by the temperature-induced methylation changes. Investigations of how the proteome is affected by these changes could be a future direction of investigations.

Our data confirmed that the presence of REs has a positional effect on gene expression, as genes with nearby REs (that is, closer than 2 kb) had significantly different expression levels. The general effect of nearby REs was a reduction of gene expression, in line with previous reports (Hollister & Gaut 2009; Wang, Weigel & Smith 2013; Hirsch & Springer 2017). Interestingly, the increased RE methylation induced by the elevated temperature treatment caused both up- and downregulation of nearby genes in our study, showing that the impact of increased methylation of nearby RE does not necessarily result in downregulation of a nearby gene.

Only four DEDMGs occured in all ecotypes, and interestingly the co-occurring methylation and gene expression was observed for the same DNA methylation context and in the same genomic locations in all or most ecotypes observed. The similar location of the methylation response in all ecotypes suggests a mechanism that recognises the same genic regions and induces DNA methylation upon temperature treatment. A similar correlation of methylome and gene expression between ecotypes has been described previously. For example, 20% of 1500 to 2500 protein coding genes that were correlated for gene expression and DNA methylation where shared between to ecotypes of lotus (Nelumbo nucifera) (Li, Yang, Wang, Chen & Shi 2021b).

### DEDMGs and observed plant phenotypes

A general observation from our DNA methylation and gene expression analysis is that each ecotype had its own set of differentially methylated regions (DMRs), differentially methylated genes (DMGs), and both differentially expressed genes and differentially methylated genes (DEDMGs) in response to temperature treatment.

NOR2 and ICE2 had several DEDMGs associated with flowering. ICE2 had a PEBP gene family FLOWERING LOCUS T (FT) homologous gene, the MADS-box transcription factors AGAMOUS (AG) and AGAMOUS-LIKE-24 (AGL24), VERNALIZATION 1 (VRN1), and *ELONGATED HYPOCOTYL 5 (HY5)* DEDMGs. NOR2 had *PHYTOCHROME AND FLOWERING TIME REGULATORY PROTEIN (PTF1)* and *HY5*. Even though flowering time was not significantly affected by temperature treatment in ES12 and IT4, two MADS-box transcription factors *SHORT VEGETATIVE PHASE (SVP)* and a SAWADEE involved in flowering were differentially methylated and expressed in ES12 and IT4, respectively (Gregis, Sessa, Colombo & Kater 2006; Law, Vashisht, Wohlschlegel & Jacobsen 2011). As these two ecotypes hardly flowered at all, it is possible that flowering is impacted by these DEDMGs under natural conditions, where IT4 and ES12 perhaps would flower more readily than in our controlled indoor setup.

Some of the identified DEDMGs in ICE2 and NOR2 may be involved in the epigenetic memory effect we observed on flowering time in these ecotypes, although more detailed gene expression analysis during the flower-induction treatment would be needed to clarify their potential impact. Previous studies have identified the *FT* homolog *FvFT1* as an important regulator of flowering time in *F. vesca*, with a hypothesized function as a floral repressor in seasonal flowering ecotypes (Koskela *et al.* 2012; Mouhu *et al.* 2013; Kurokura *et al.* 2017). In contrast to this hypothesis, we found that *FvFT1* was a DEDMG with >10-fold downregulation in the elevated temperature treatment and delayed flowering in the ICE2 ecotype. On the other hand, *AGL24* and *VRN1*, both known to promote flowering in *Arabidopsis* (Yu, Xu, Tan & Kumar 2002; Sung & Amasino 2004), were downregulated DEDMGs in ICE2, demonstrating a positive correlation between gene expression and the flowering phenotype we observed.

PHYTOCHROME INTERACTING FACTOR (PIF4) is reported to be a key factor in plant responses to temperature and could interact with the transcription factor HY5 (Kumar *et al.* 2012; Lee, Wang & Huq 2021). HY5, together with HISTONE DEACETYLASE 9 (HDA9), has been reported to directly bind to the promoter region of PIF4, a crucial regulator of plant thermomorphogenesis and a flowering regulator CONSTANS LIKE 5 (COL5) that promotes flowering in short day-grown *Arabidopsis* (Hassidim, Harir, Yakir, Kron & Green 2009; Chu *et al.* 2022). The observed >3-fold reduction in HY5 transcripts following the elevated temperature treatment therefore fits with the delayed flowering phenotype observed in ICE2 and NOR2. Thus, the differential methylation and expression of HY5 could at least partly explain why NOR2 and ICE2 had a more plastic flowering response to increased temperature than the other ecotypes.

### Temperature-induced methylation of the epigenetic machinery

Twenty-one of the DEDMGs we identified are themselves part of the DNA methylation machinery, either directly or indirectly through effects on other chromatin modifications. None of these epigenetic-related DEDMGs were shared by all ecotypes - in fact only four such DEDMGs were shared and then only by two ecotypes. A change of expression of these proteins due to DNA methylation induced by the temperature treatments may ultimately lead to further chromatin changes in an ecotype-specific manner. Several examples of DEDMGs with roles in the DNA methylation machinery are already discussed in this report.

These include the aforementioned SAWADEE homolog involved in RNA-directed DNA methylation and a small RNA-degrading nuclease (Law *et al.* 2011, 2013). Another example is a Jumonji domain-containing (Jmjc) histone demethylases such as the JMJ20, that may reinforce histone H3K9me2 and enhance CMT3/2 CHG/CHH methylation (Miura *et al.* 2009; Greenberg *et al.* 2013).

We have in this study demonstrated a clear memory effect on the phenotypic level that is accompanied by DNA methylation and transcriptional changes. We have also shown that the effect is ecotype (genotype) specific and that some ecotypes are more plastic than others. This lays the foundation for new avenues of investigations, also including sRNA and histone modifications, to decipher how the environment sets its marks on DNA.

## Supplementary figure legends

**Fig S1. Experimental design: propagation, sample collection, and phenotypic observations of asexually propagated *Fragaria vesca* plants**. Four ecotypes of *F. vesca* were propagated for three asexual generations (AS1-3) through stolon formation. Plants were propagated under normal (18 °C; blue boxes) and elevated temperature conditions 28 °C; red boxes). Phenotypic observations were made in a common garden environment (green boxes).

**Fig S2. Stolon production in *Fragaria vesca* ecotypes propagated at normal (18 ºC) and elevated (28 ºC) temperature conditions.** The number of stolons produced per plant was recorded bi-weekly for 14 weeks after short day treatment of plants propagated for one (left panels) or three asexual generations (right panels) at 18 °C (blue lines) or 28 °C (red lines).

**Fig S3. DNA methylation landscape in leaves of *Fragaria vesca* ecotypes ES12, ICE2, and IT4 propagated under normal temperature conditions (18 ºC).** Violin plots showing the DNA methylation level (methylated reads/total reads) of different methylation contexts on all seven *F. vesca* (Fv) chromosomes. The 25 quantile and 75 quantile values are indicated by the lower and upper margins of the boxplot and the median is indicated by the black line between quantiles. Error bars show 95% confidence intervals. Ecotypes shown are ES12, ICE2, and IT4.

**Fig S4. DNA methylation differences in leaves of *Fragaria vesca* ecotypes propagated at normal (18 ºC) temperature conditions (AS3).** Plots show methylation level (methylated reads/total reads) in ecotypes ES12, ICE2, IT4 and NOR2). The methylation level is shown for all methylation contexts along all seven *F. vesca* (Fv) chromosomes, using a 50 kb window.

**Fig S5. DNA methylation differences in leaves of *Fragaria vesca* ecotypes, using ecotype ES12 as a baseline.** Plots show differences in methylation level (methylated reads/total reads) between ecotype ICE2, IT4, and NOR2 propagated at 18 ºC for three asexual generations. Methylation differences are shown for different methylation contexts (CGN, CHG, CHH) along all seven *F. vesca* (Fv) chromosomes using ES12 as a reference, using a 50 kb window.

**Fig S6. CGN DNA methylation in leaves of *Fragaria vesca* ecotypes propagated at normal (18 ºC) temperature conditions showing CGN trinucleotide context.** The plots show CGN methylation level (methylated reads/total reads) in different ecotypes (ES12, ICE2, IT4, NOR2) along all seven *F. vesca* (Fv) chromosomes for all CGN trinucleotide contexts (CGA, CGC, CGG, CGT) using a 50kb window.

**Fig S7. CHG DNA methylation in leaves of *Fragaria vesca* ecotypes propagated at normal (18 ºC) temperature conditions showing CHG trinucleotide context.** The plots show CHG methylation level (methylated reads/total reads) in different ecotypes (ES12, ICE2, IT4, NOR2) along all seven *F. vesca* (Fv) chromosomes for all CHG trinucleotide contexts (CAG, CCG, CTG) using a 50kb window.

Fig S8. CHH DNA methylation in leaves of *Fragaria vesca* ecotypes propagated at normal (18 ºC) temperature conditions showing CHH trinucleotide context. The plots show CHH methylation level (methylated reads/total reads) in different ecotypes (ES12, ICE2, IT4, NOR2) along all seven *F. vesca* (Fv) chromosomes for all CHH trinucleotide contexts (CAA, CAC, CAT, CCA, CCC, CCT, CTA, CTC, CTT) using a 50kb window.

**Fig S9. Correlation between methylation level and gene density in leaves of different *Fragaria vesca* ecotypes propagated at normal (18 ºC) temperature.** Plots show methylation level (methylated reads/total reads) and gene density along all seven *F. vesca* (Fv) chromosomes in ecotype ES12 **(A)**, ICE2 **(B)**, and IT4 **(C)**, using a 50 kb window.

**Fig S10. DNA methylation differences between *Fragaria vesca* propagated at normal (18 ºC) and elevated (28 ºC) temperature conditions (AS3).** Plots show methylation differences between temperatures (28 vs. 18 ºC) in four ecotypes (ES12, ICE2, IT4, NOR2). Differences in methylation level (methylated reads/total reads) are shown for different methylation contexts (CGN, CHG, CHH) along all seven *F. vesca* (Fv) chromosomes, using a 50 kb window.

**Fig S11. CGN DNA methylation differences between *Fragaria vesca* propagated at normal (18 ºC) and elevated (28 ºC) temperature conditions showing CGN trinucleotide context.**Plots show differences in methylation level (methylated reads/total reads) between temperatures (28 vs. 18 ºC) in different ecotypes (ES12, ICE2, IT4, NOR2) along all seven *F. vesca* (Fv) chromosomes and for all CGN trinucleotide contexts (CGA, CGC, CGG, CGT) using 18 ºC as baseline and a 50kb window.

**Fig S12. CHG DNA methylation differences between *Fragaria vesca* propagated at normal (18 ºC) and elevated (28 ºC) temperature conditions showing CHG trinucleotide context.** Plots show differences in methylation level (methylated reads/total reads) between temperatures (28 vs. 18 ºC) in different ecotypes (ES12, ICE2, IT4, NOR2) along all seven *F. vesca* (Fv) chromosomes and for all CHG trinucleotide contexts (CAG, CCG, CTG) using 18 ºC as baseline and a 50kb window.

**Fig S13. CHG DNA methylation differences between *Fragaria vesca* propagated at normal (18 ºC) and elevated (28 ºC) temperature conditions showing CHH trinucleotide context.** Plots show differences in methylation level (methylated reads/total reads) between temperatures (28 vs. 18 ºC) in different ecotypes (ES12, ICE2, IT4, NOR2) along all seven *F. vesca* (Fv) chromosomes and for all CHH trinucleotide contexts (CAA, CAC, CAT, CCA, CCC, CCT, CTA, CTC, CTT) using 18 ºC as baseline and a 50kb window.

**Fig S14. DNA methylation patterns of protein-coding genes, repetitive elements (REs), and pseudogenes in leaves of different *Fragaria vesca* ecotypes propagated at normal (18 ºC) and elevated (28 ºC) temperature conditions for three asexual generations** Plots show methylation levels in protein-coding genes **(A,D,G)**, REs **(B,E,H)**, and pseudogenes **(C,F,I)** in ecotype ES12 **(A-C)**, ICE2 **(D-F)**, and IT4 **(G-I)**. Each genomic feature, and regions 2 kb up- and downstream to these, were divided into 20 pieces to calculate methylation levels (methylated reads/total reads). Colored lines show different combinations of temperature conditions and methylation contexts (CGN, CHG, CHH).

**Fig S15. DNA methylation landscape in leaves of *Fragaria vesca* plants propagated at normal (18 ºC) temperature conditions for one to three generations.** Plots show methylation differences between different ecotypes (ES12, ICE2, IT4, NOR2) propagated for three (AS3) vs. one (AS1) asexual generation. Methylation levels (methylated reads/total reads) along each *F. vesca* (Fv) chromosome are shown for different methylation contexts (CGN, CHG, CHH), using AS1 as a baseline and a 50 kb window.

**Fig S16. Chromosomal and genic distribution patterns of differentially methylated regions (DMRs) in leaves of *Fragaria vesca* plants propagated at normal (18 ºC) temperature conditions for one to three generations.** Circos plots show patterns of DMRs between plants propagated for one vs. three asexual generations for all seven *F. vesca* chromosomes in ecotype ES12 **(A)**, ICE2 **(B)**, IT4 **(C)**, and NOR2 **(D)**. Lanes 1-7 show gene density, DMR density and methylation level (%) for different methylation contexts (CGN, CHG, and CHH). Connecting lines in the middle indicate coding regions (CDS) regions that showed synteny.

**Fig S17. Chromosomal and genic distribution patterns of differentially methylated regions (DMRs) in leaves of *Fragaria vesca* plants propagated at normal (18 ºC) and elevated (28 ºC) temperature conditions for three asexual generations.** Circos plots show patterns of DMRs between plants propagated at 28 vs. 18 ºC for all seven *F. vesca* (Fv) chromosomes in ecotype ES12 (A), ICE2 (B), and IT4 (C). Lanes 1-7 show gene density, DMR density and methylation level (%) for different methylation contexts (CGN, CHG, and CHH). Connecting lines in the middle indicate coding regions (CDS) regions that showed synteny.

**Fig S18. Volcano plots showing differentially expressed genes (DEGs) in leaves of *Fragaria vesca* ecotypes propagated at normal (18 ºC) and elevated (28 ºC) temperature conditions.** Plots show DEGs between 28 ºC vs. 18 ºC in ecotype ES12 **(A)**, ICE2 **(B)**, and IT4 **(C)**. Blue and red dots show down- and upregulated genes, respectively. Dashed lines delineate P-value ≤ 0.05 and log_2_FoldChange ≥ 1.5.

**Fig S19. DNA methylation and gene expression change in differentially expressed and differentially methylated genes (DEDMGs) in the ‘epigenetics’ function category in leaves of different *Fragaria vesca* ecotypes (ES12, ICE2, IT4, NOR2) propagated at normal (18 ºC) and elevated (28 ºC) temperature conditions.** DEDMGs that are shared between ecotypes are highlighted in yellow.

**Fig S20. DNA methylation and gene expression change in differentially expressed and differentially methylated genes (DEDMGs) in the ‘flowering-related’ function category in leaves of different *Fragaria vesca* ecotypes (ES12, ICE2, IT4, NOR2) propagated at normal (18 ºC) and elevated (28 ºC) temperature conditions. All the identified DEDMGs were on *F. vesca* chromosome 4 (FvH4).** DEDMGs that are shared between ecotypes are highlighted in yellow.

**Fig S21. DNA methylation and gene expression change in differentially expressed and differentially methylated genes (DEDMGs) in the ‘temperature-related’ function category in leaves of different *Fragaria vesca* ecotypes (ES12, ICE2, IT4, NOR2) propagated at normal (18 ºC) and elevated (28 ºC) temperature conditions.** DEDMGs that are shared between ecotypes are highlighted in yellow.

**Fig S22. DNA methylation and gene expression change in differentially expressed and differentially methylated genes (DEDMGs) in the ‘hormone-related’ function category in leaves of different *Fragaria vesca* ecotypes (ES12, ICE2, IT4, NOR2) propagated at normal (18 ºC) and elevated (28 ºC) temperature conditions.** DEDMGs that are shared between ecotypes are highlighted in yellow.

**Fig S23. Positional effect of repetitive elements (REs) on gene expression level and absolute gene expression change in leaves of the ES12, ICE2, and IT4 *Fragaria vesca* ecotypes propagated at normal (18 ºC) and elevated (28 ºC) temperature conditions.** Positional effect of REs on gene expression level **(A)** and absolute gene expression change **(B)** in indicated ecotypes. The distance from the gene to the nearest RE is given on the x-axis. The distance (in Kbp) from a gene to the nearest RE is given on the x-axis. Asterisks indicate significant differences calculated by the games_howell_test: ** 0.001 ≤ p < 0.01; *** 0.0001 ≤ p < 0.001; **** 0.00001 ≤ p < 0.0001; ns = not significant. Error bars show 95% confidence intervals.

