## Supplementary figures and images for "Temperature-induced methylome changes during asexual reproduction trigger transcriptomic and phenotypic changes in *Fragaria vesca*"

### Supplemental Figure 1

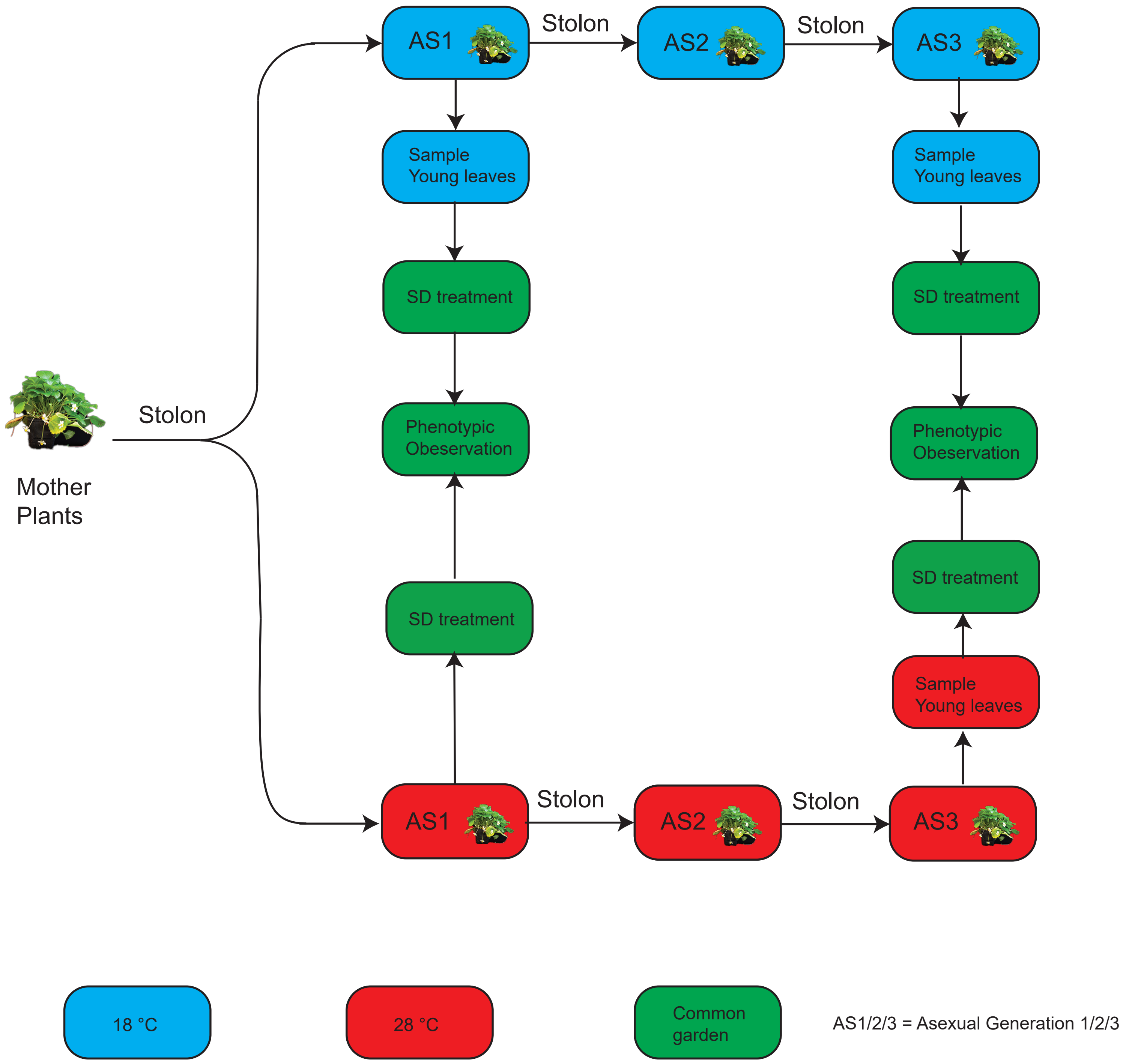

### Supplemental Figure 2

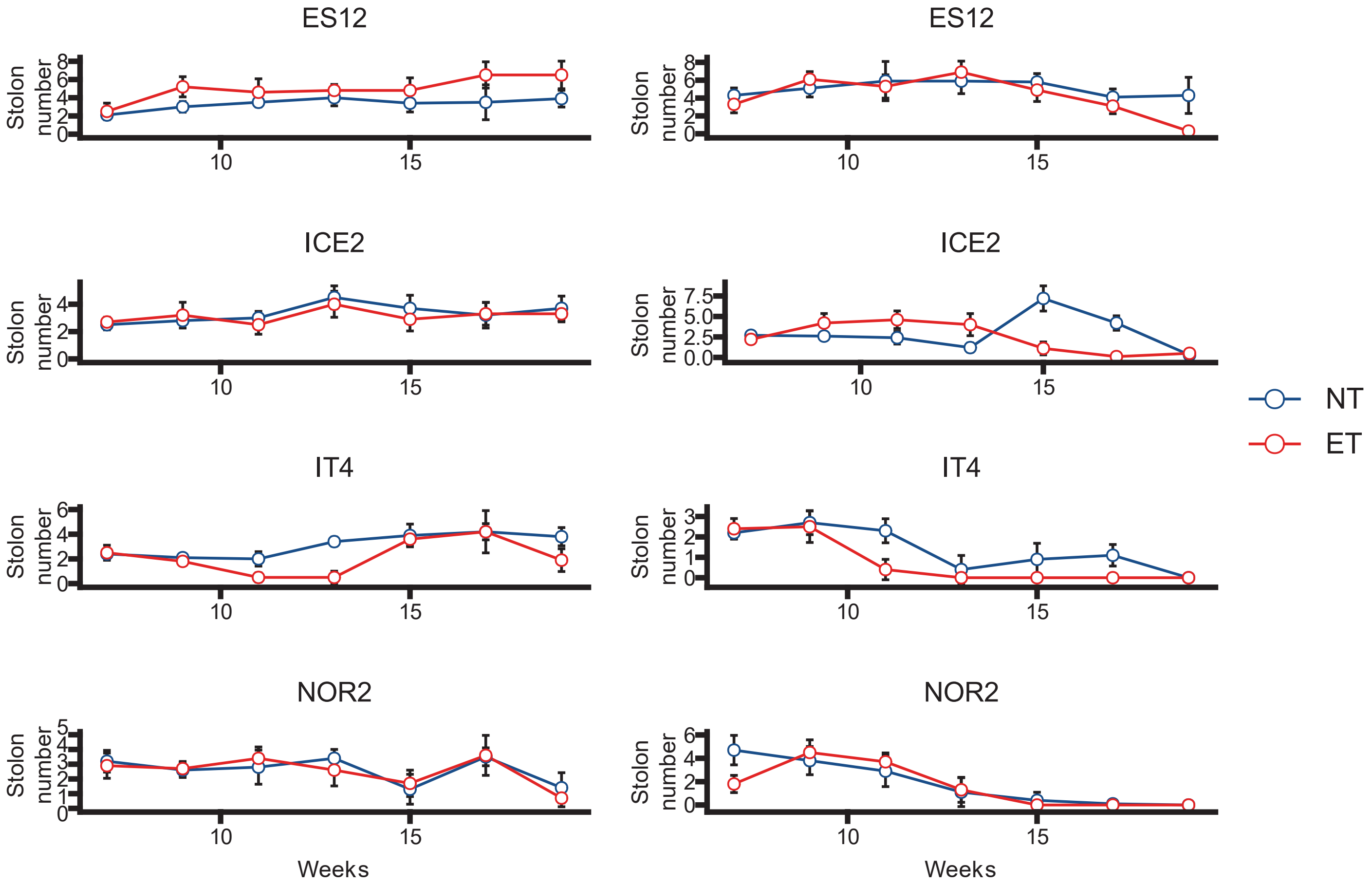


Stolon numbers of AS1

Stolon numbers of AS3

### Supplemental Figure 3

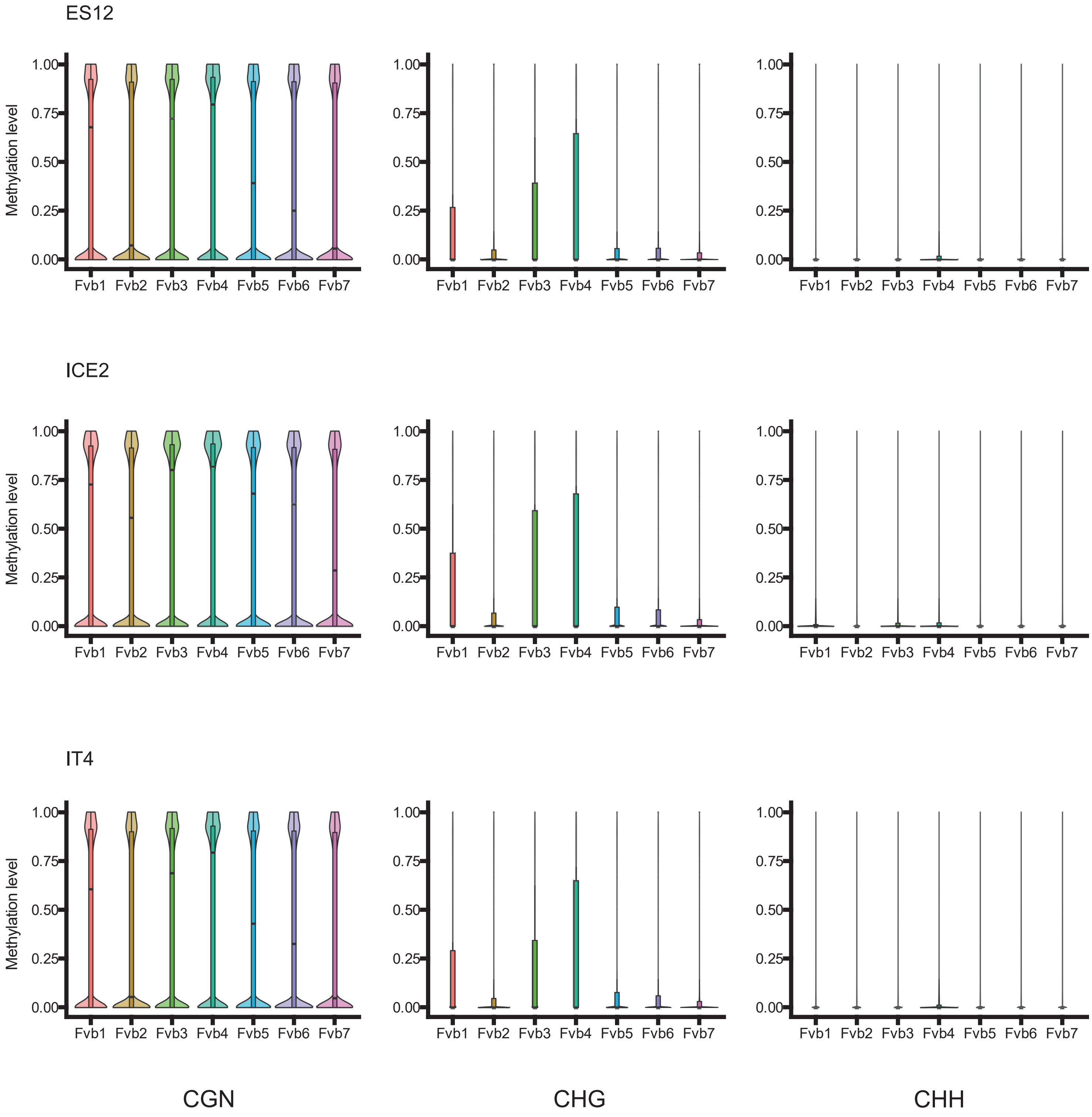

### Supplemental Figure 4

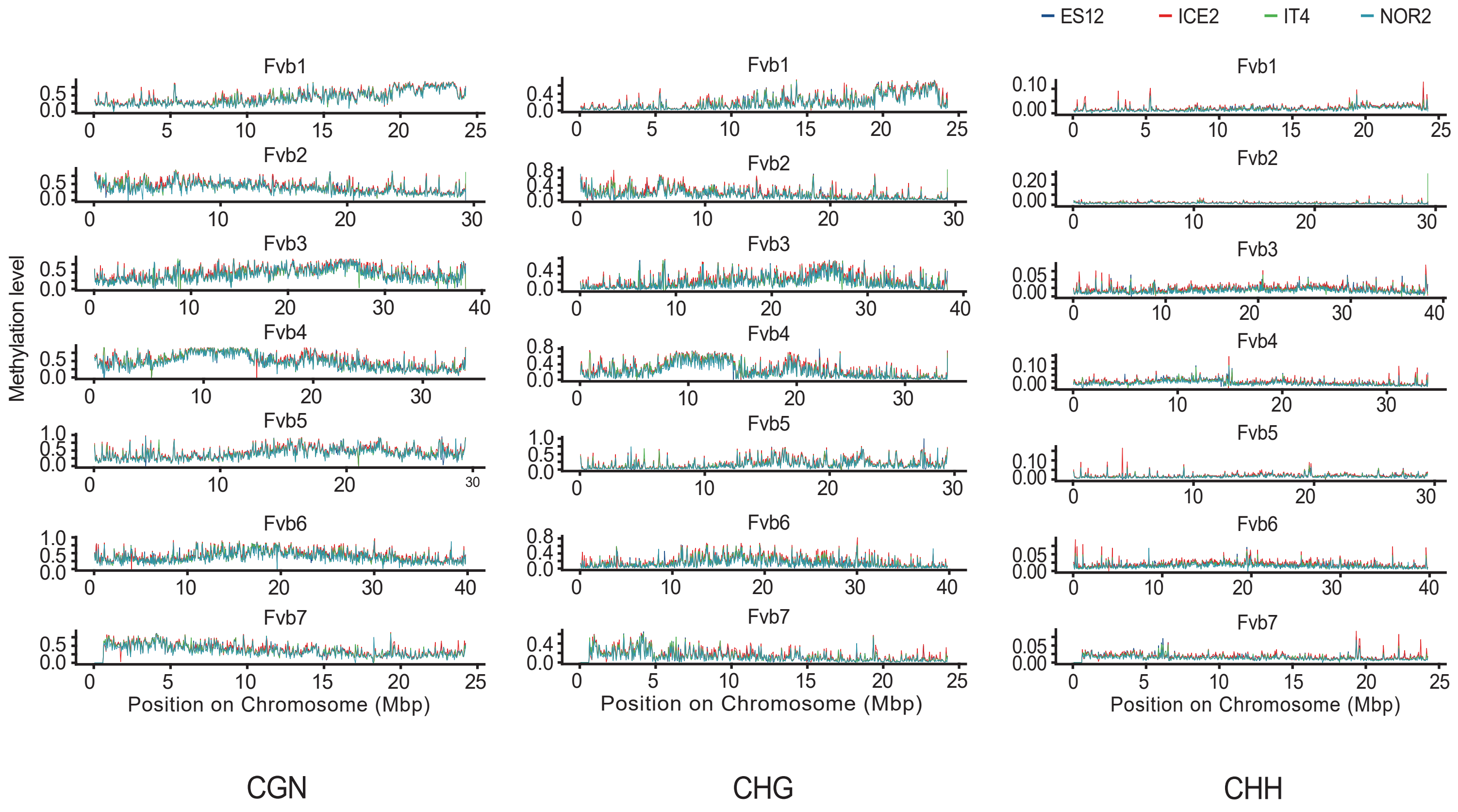

### Supplemental Figure 5

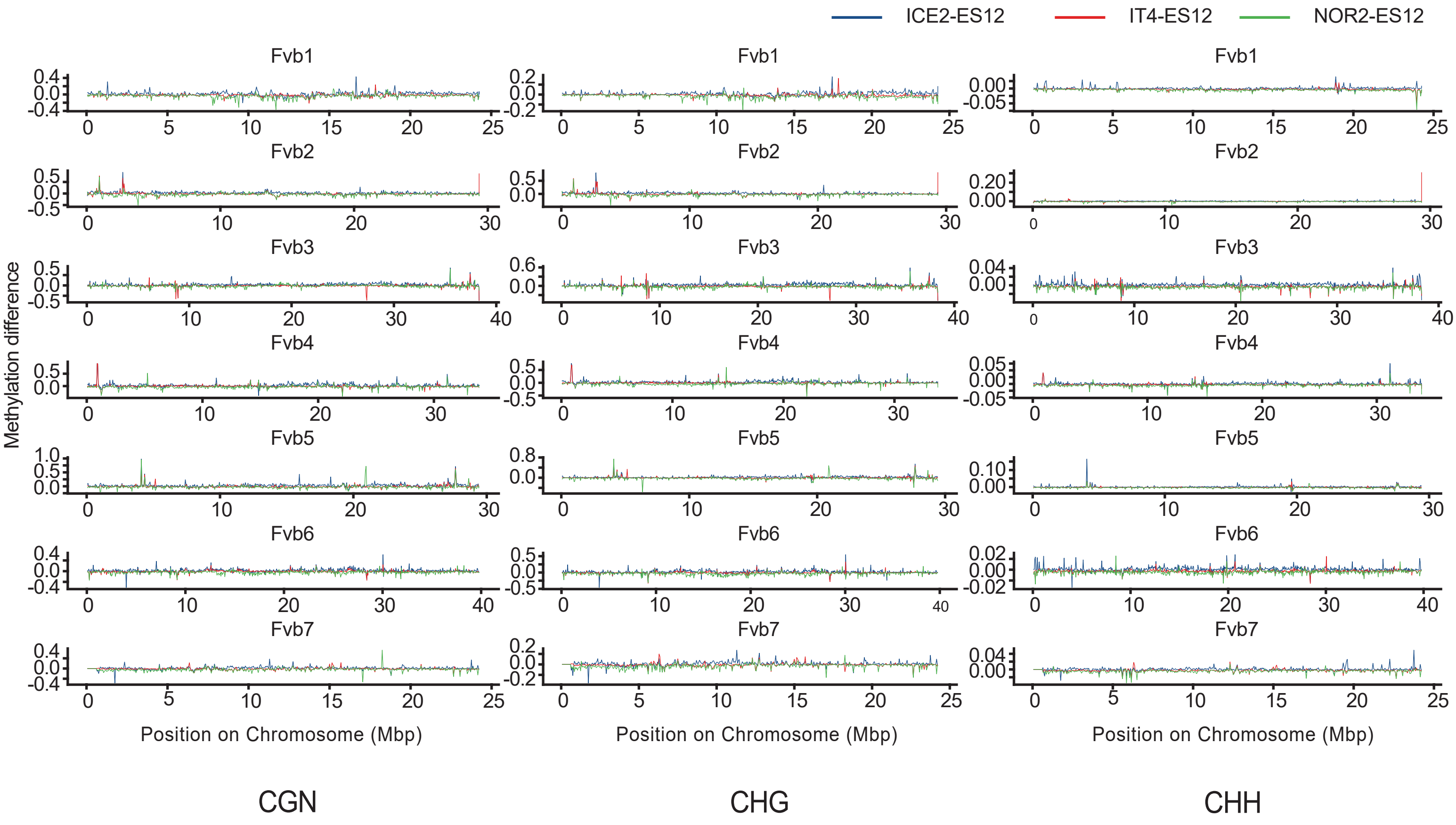

### Supplemental Figure 6

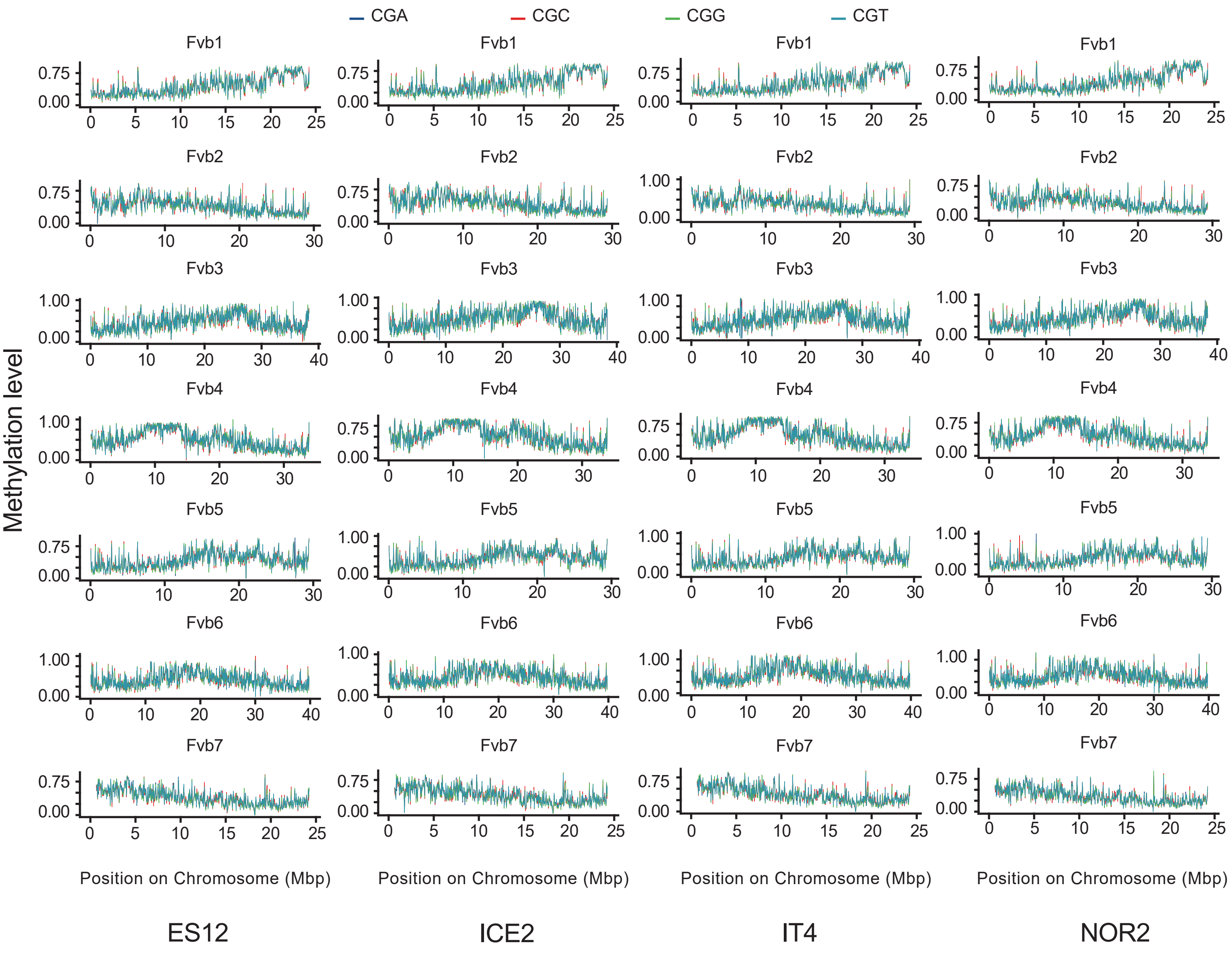

### Supplemental Figure 7

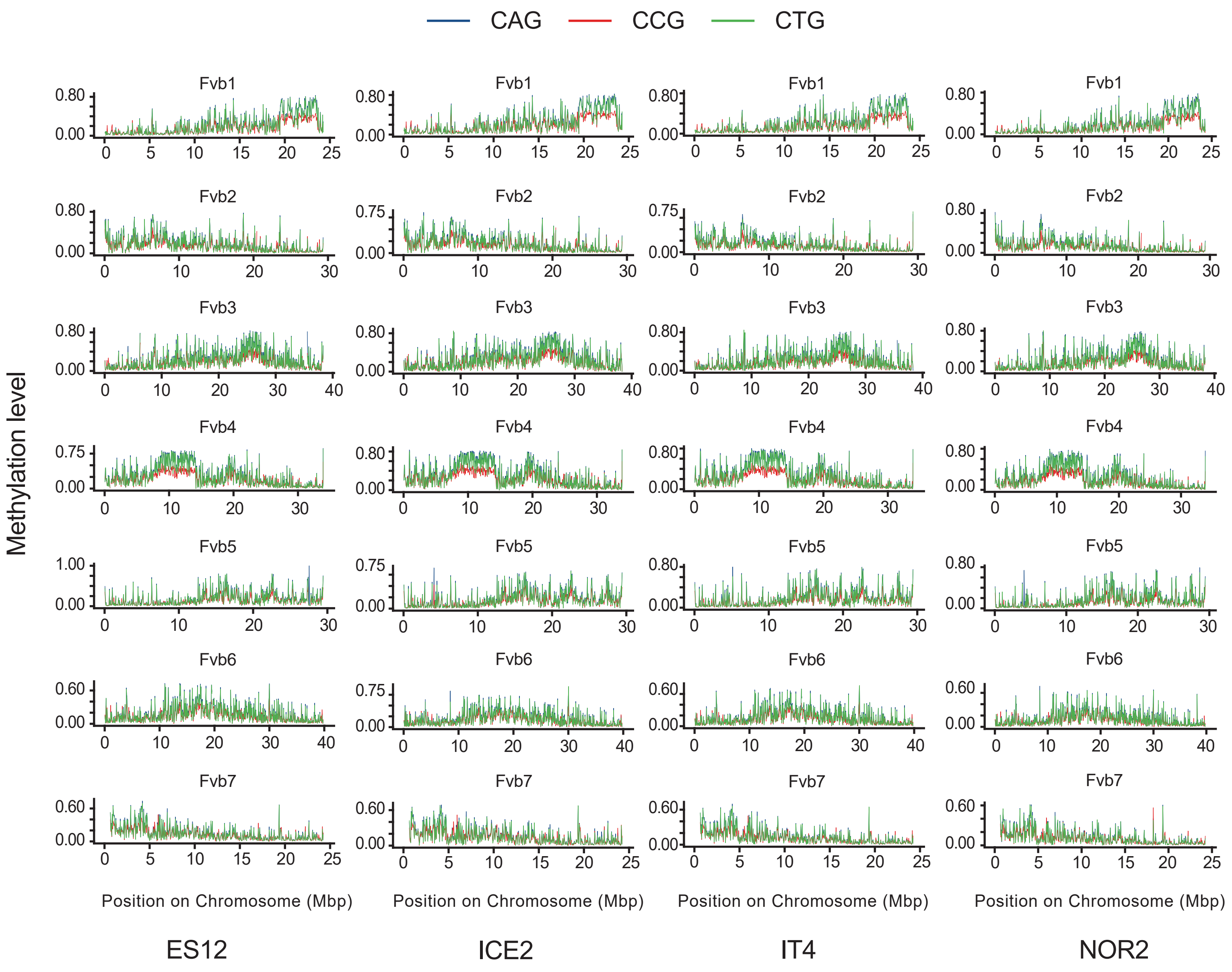

### Supplemental Figure 8

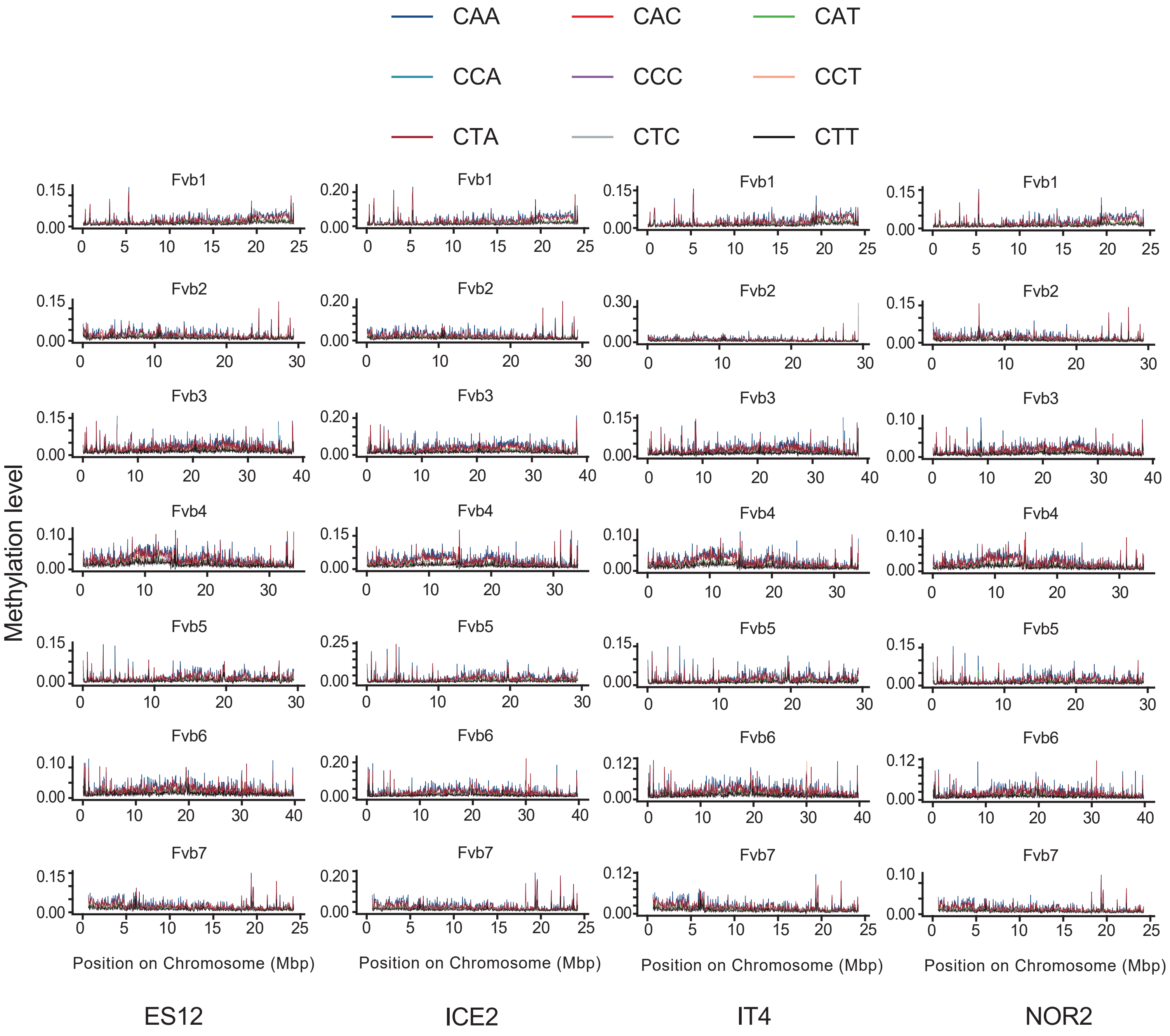

### Supplemental Figure 9

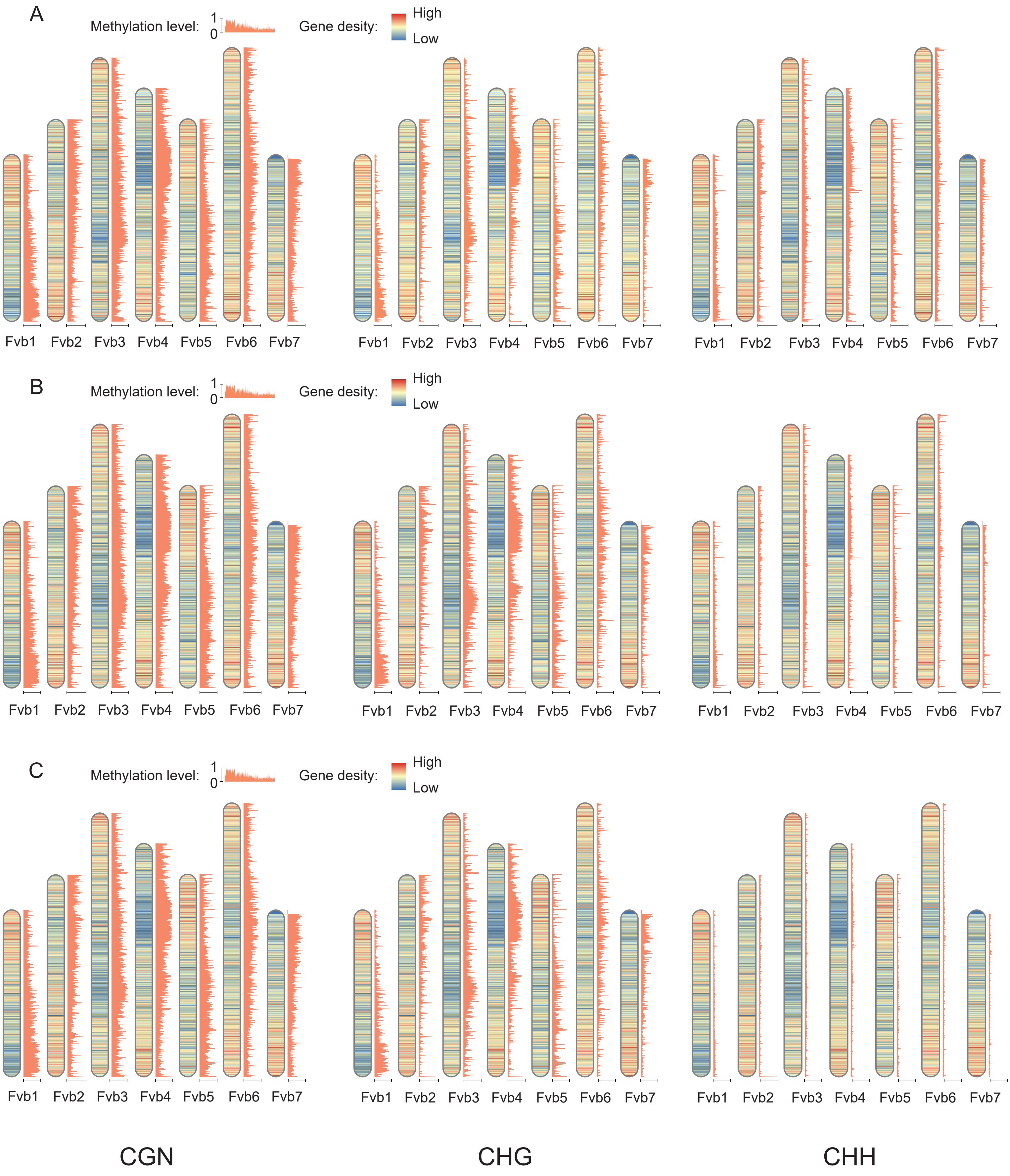

### Supplemental Figure 10

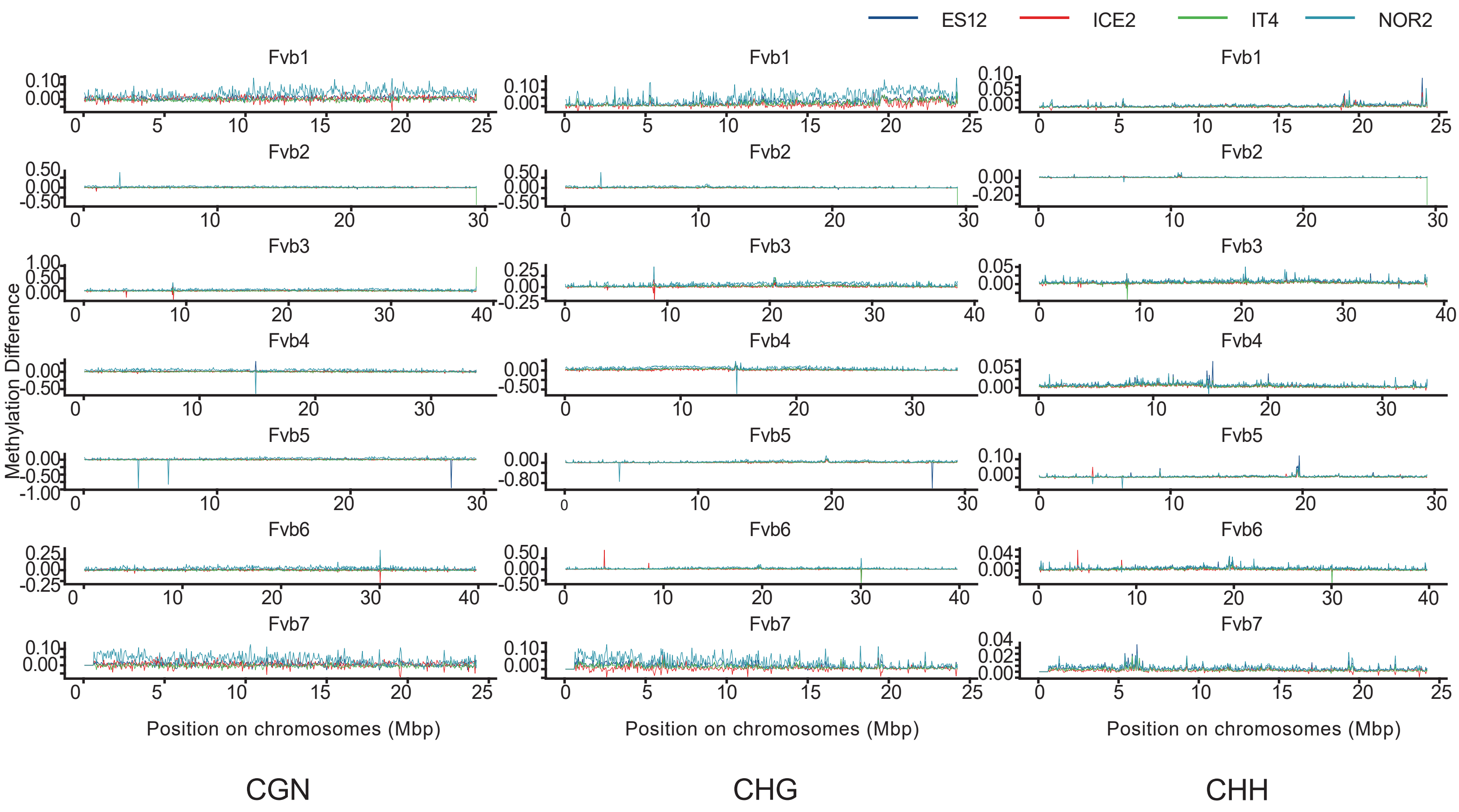

### Supplemental Figure 11

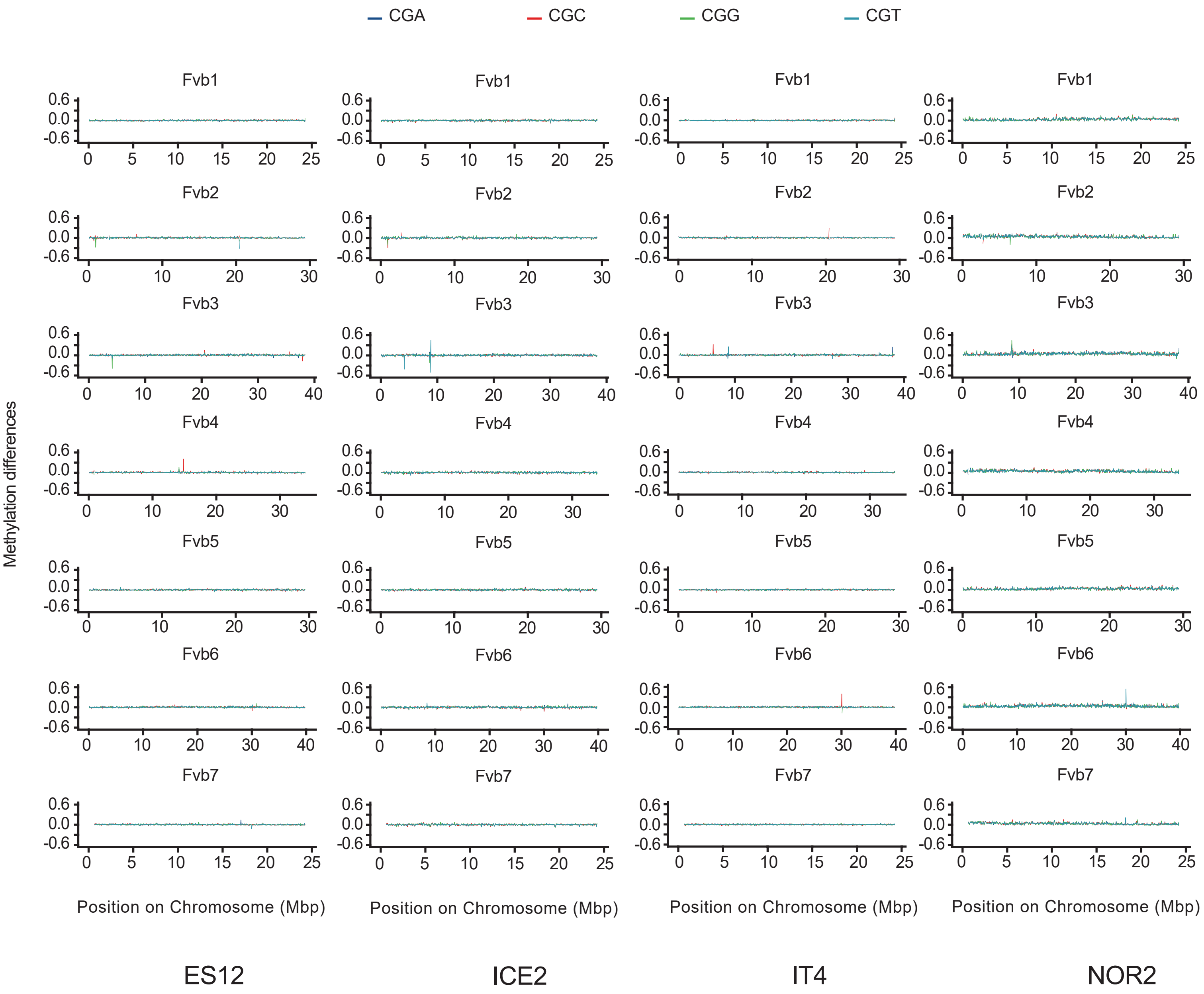

### Supplemental Figure 12

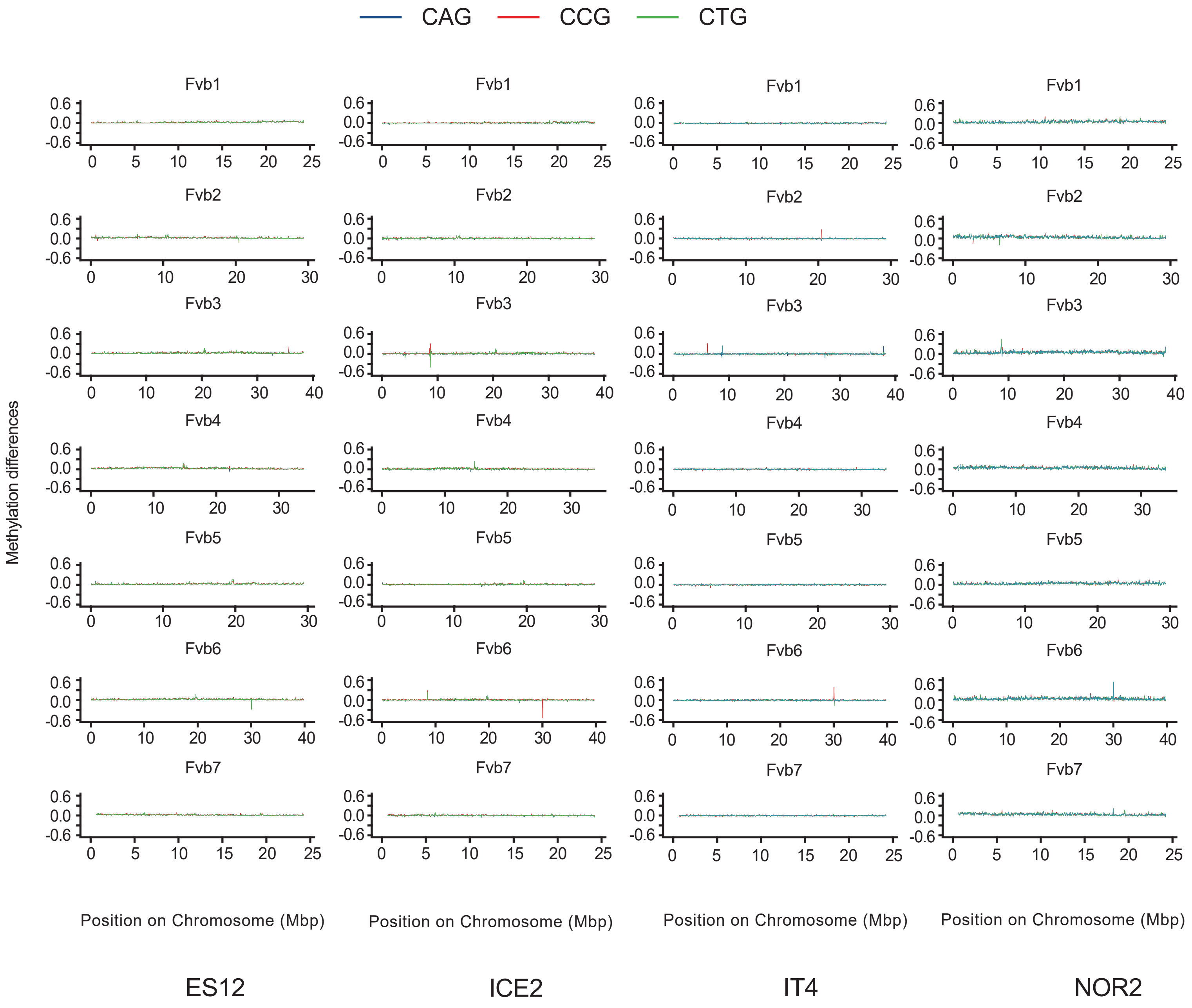

### Supplemental Figure 13

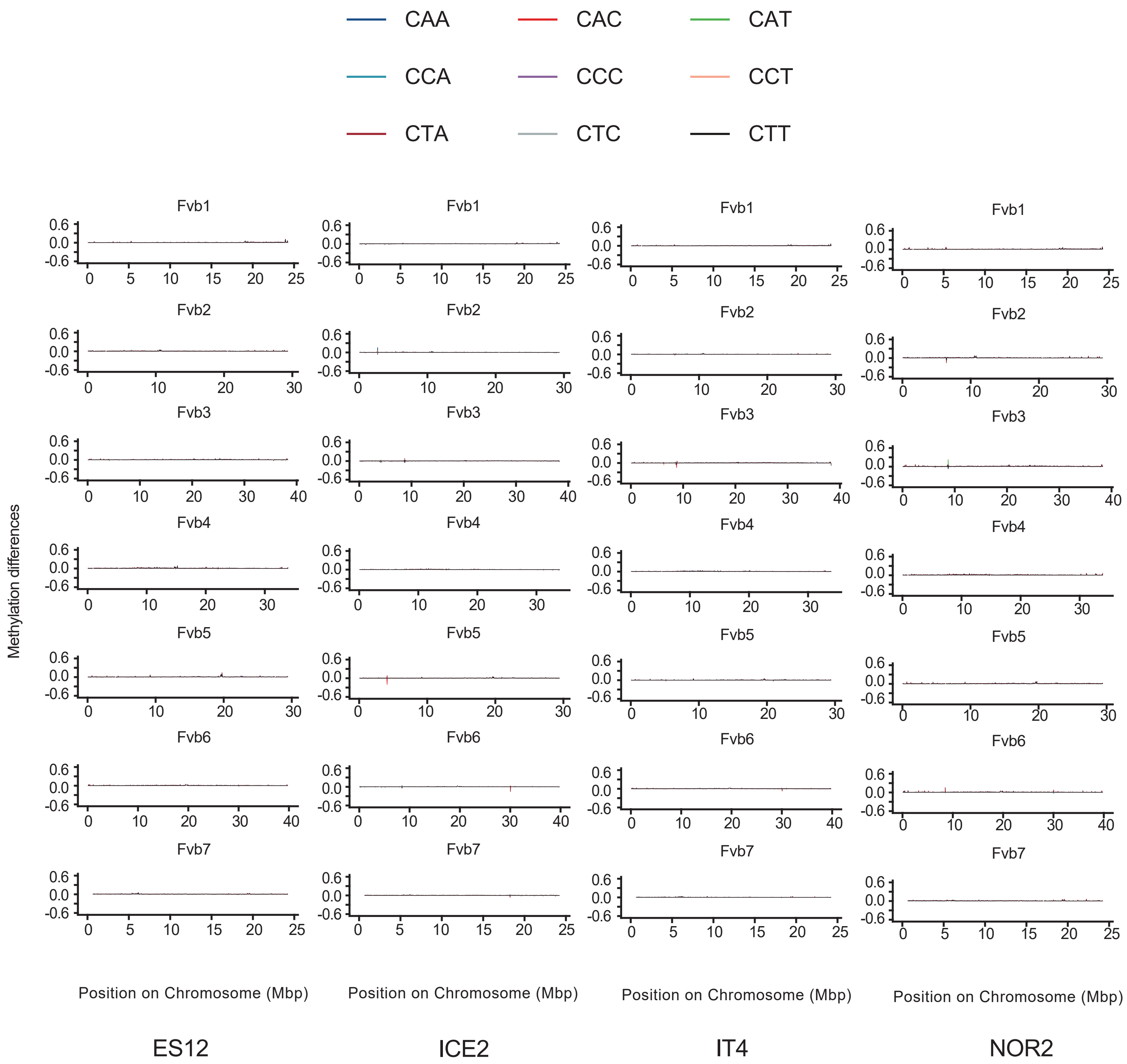

### Supplemental Figure 14

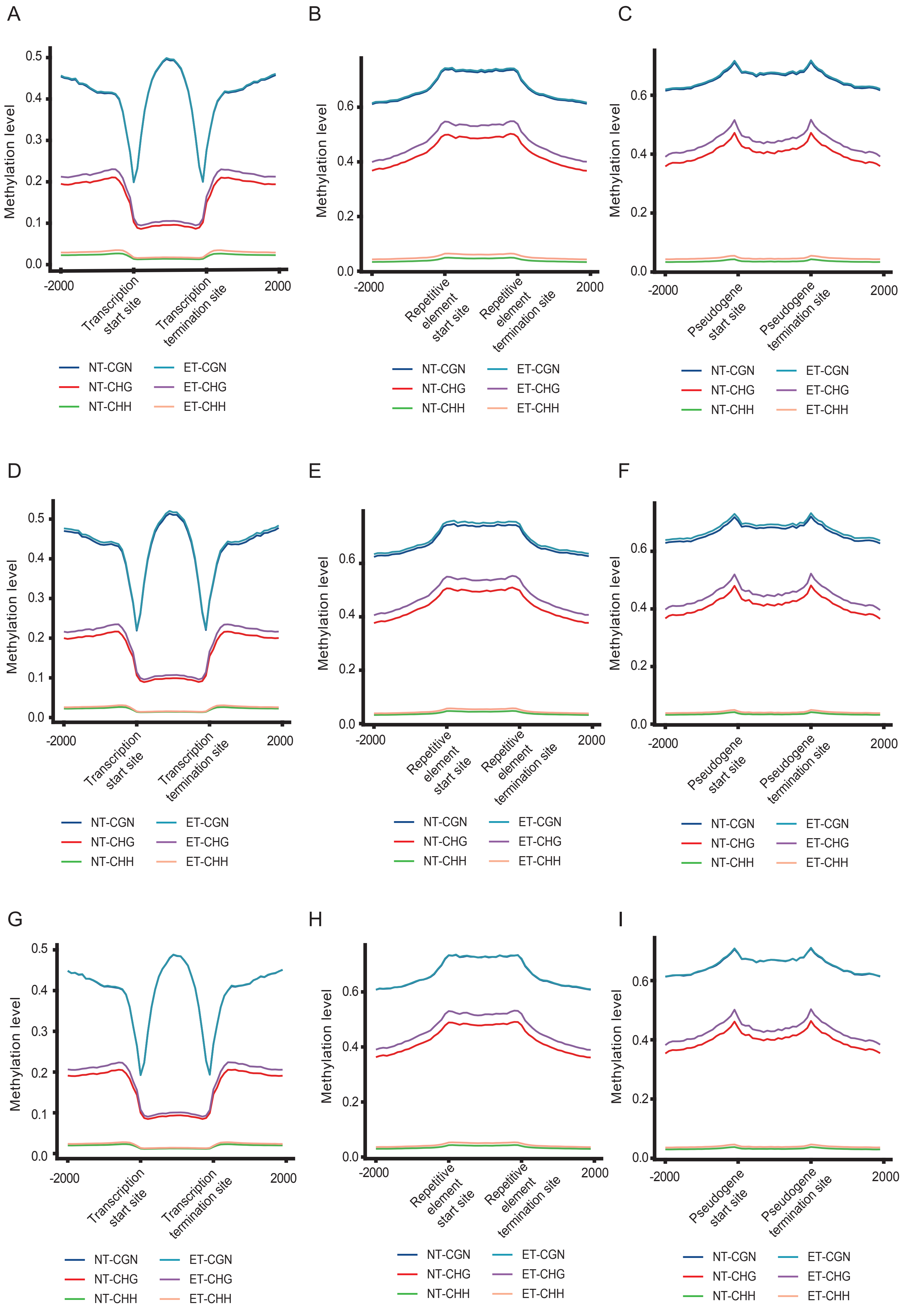

### Supplemental Figure 15

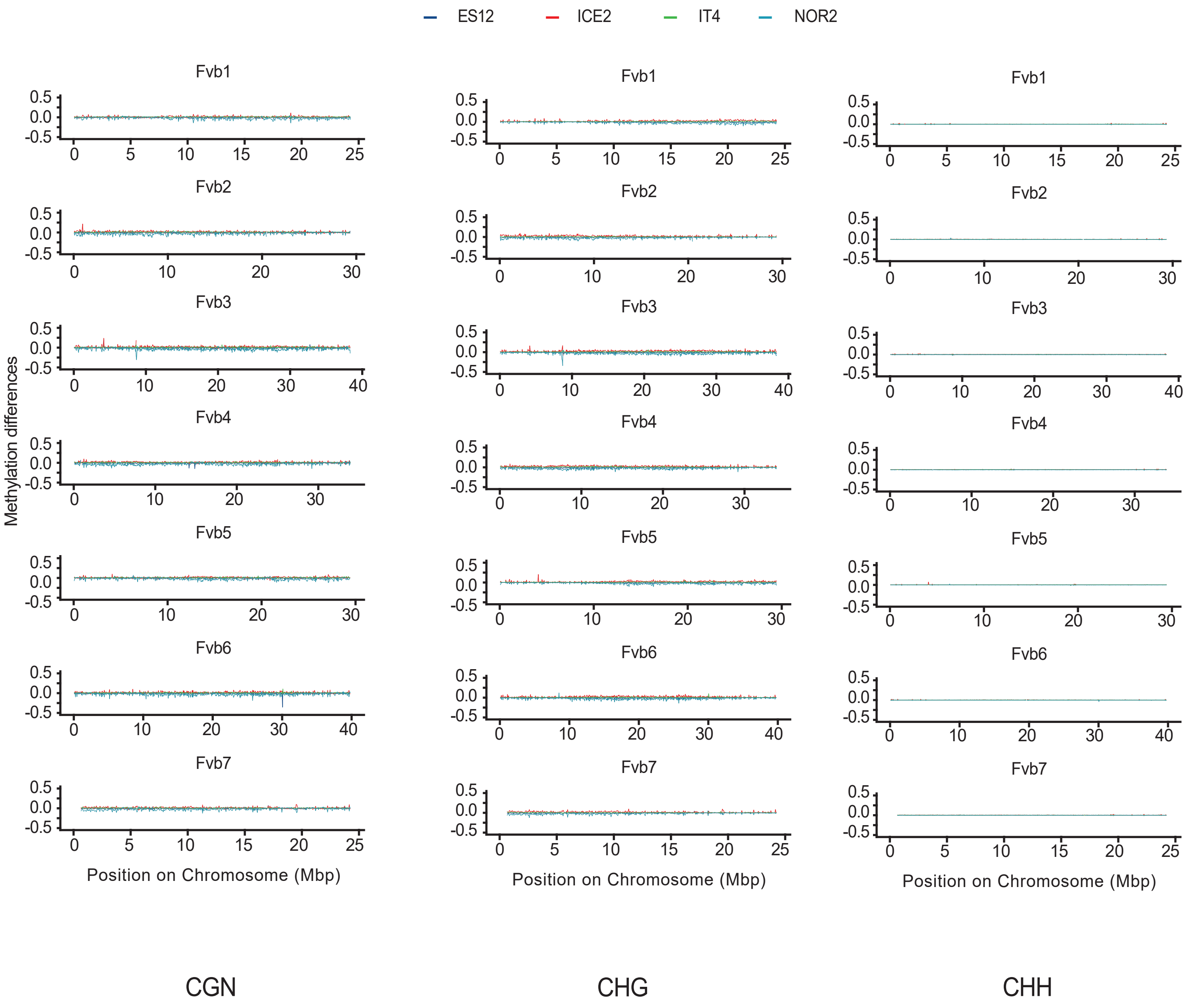

### Supplemental Figure 16

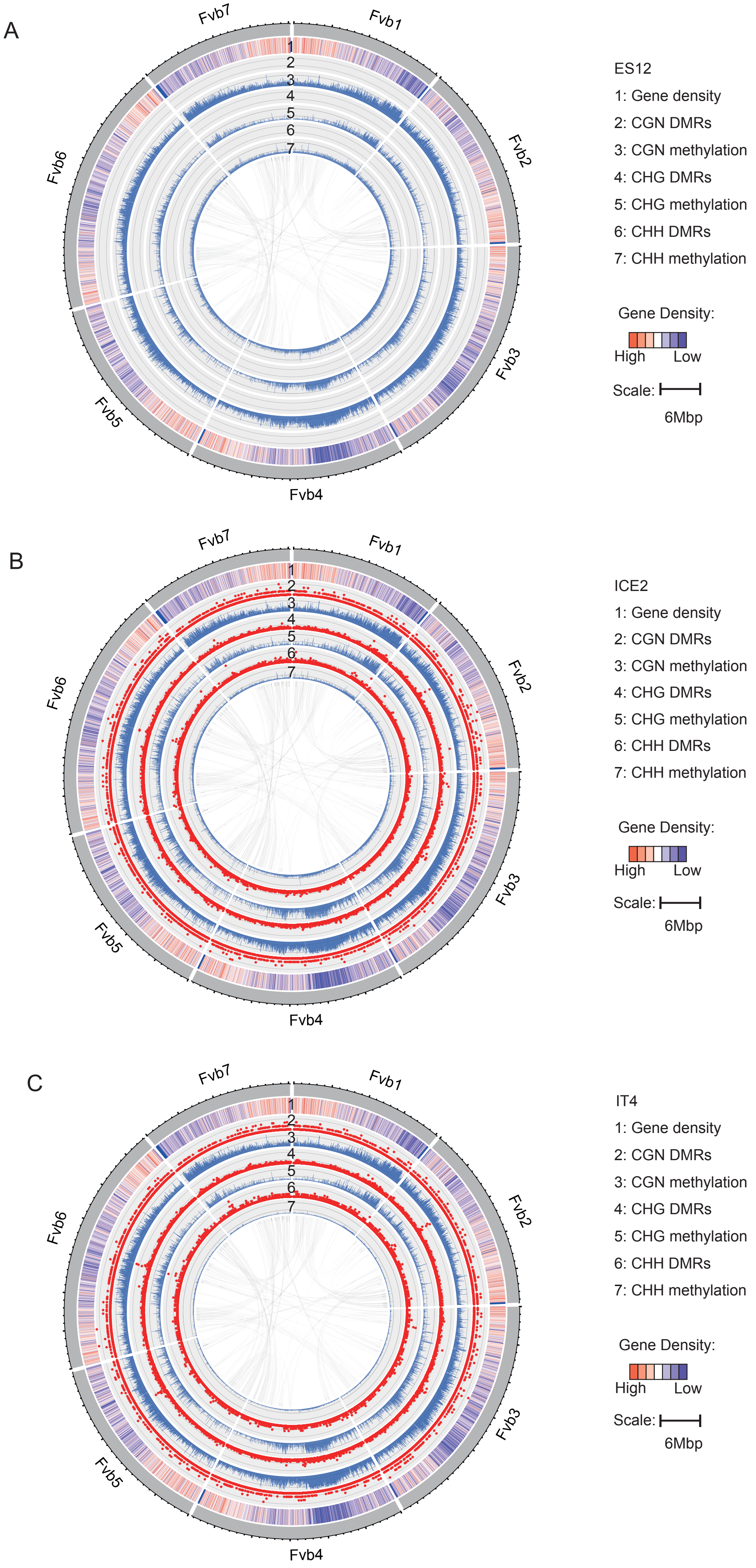

### Supplemental Figure 18

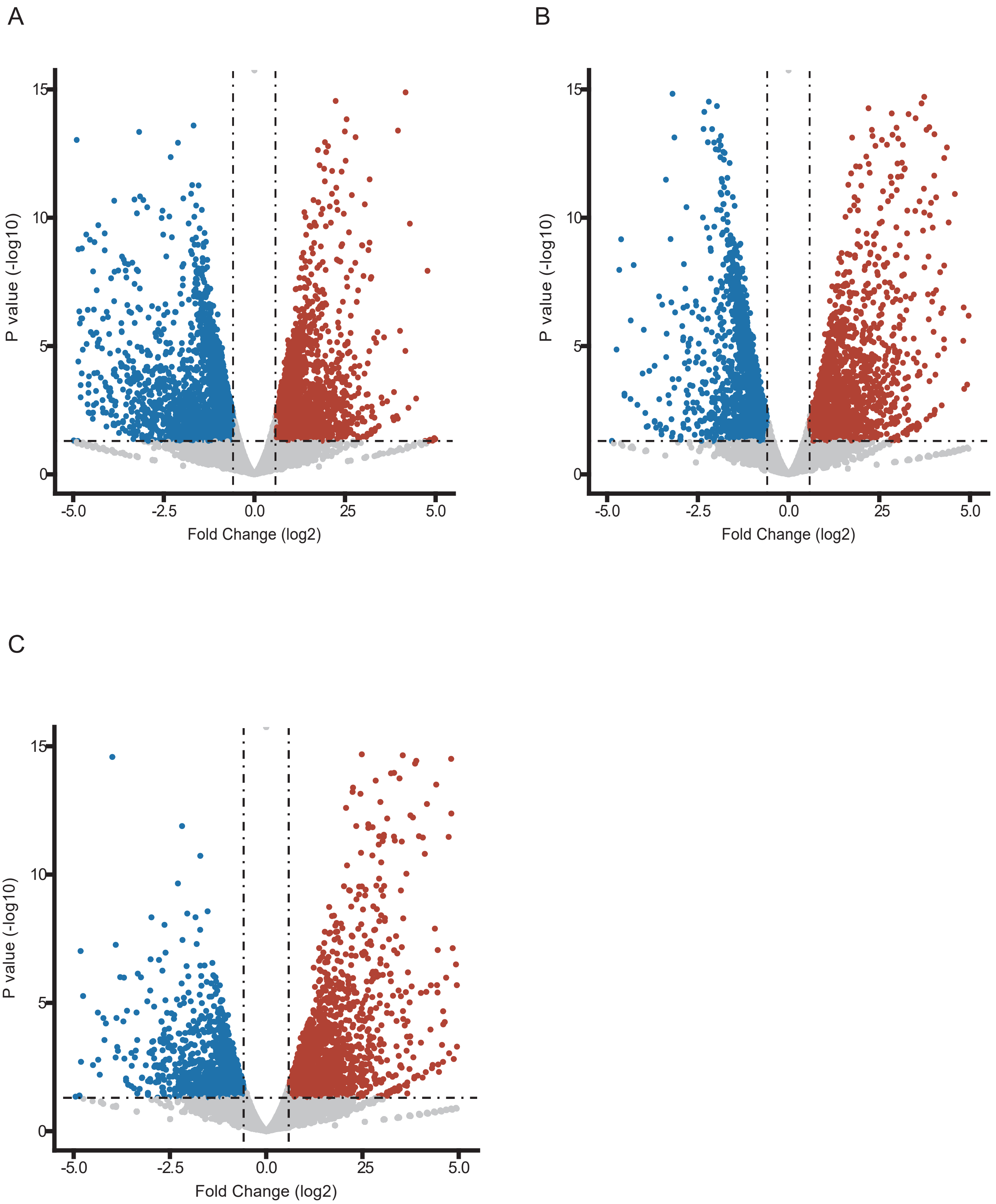

### Supplemental Figure 19

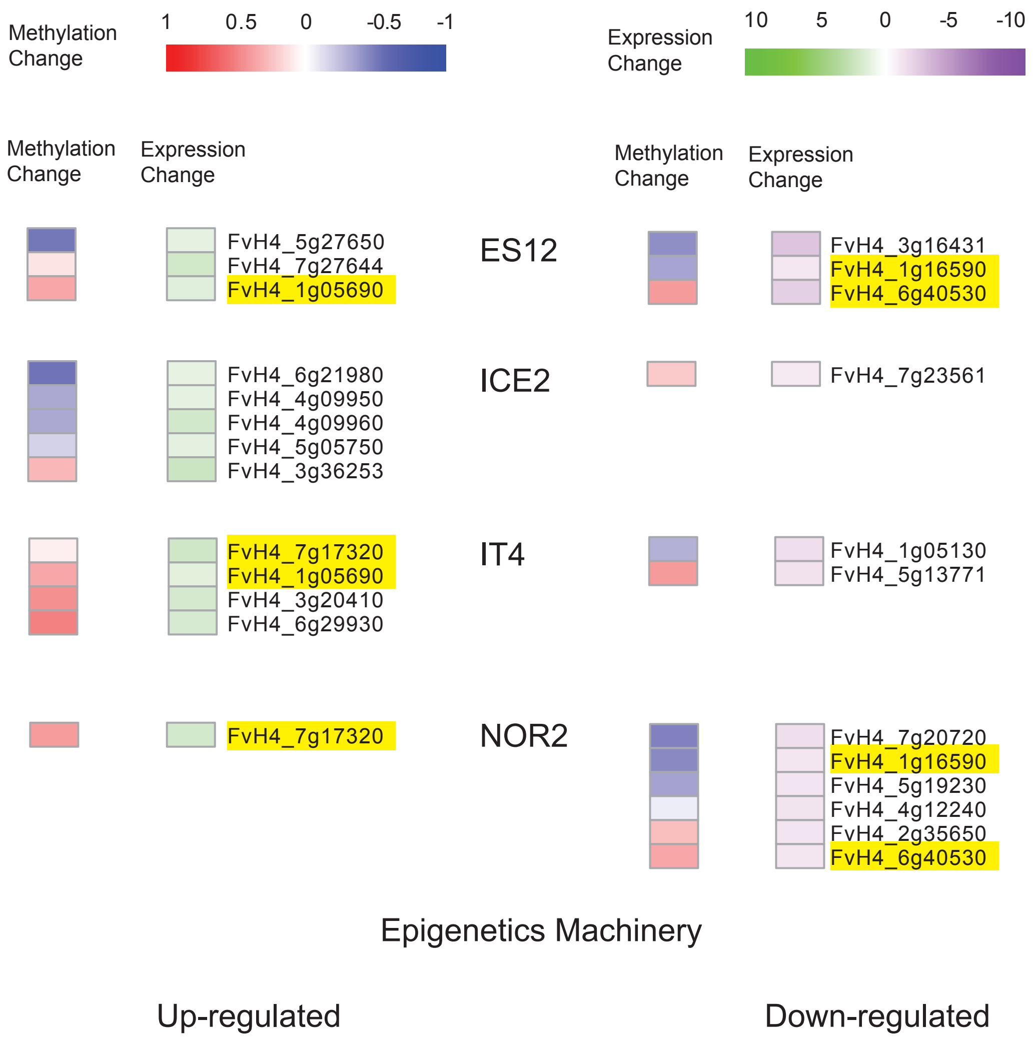

### Supplemental Figure 20

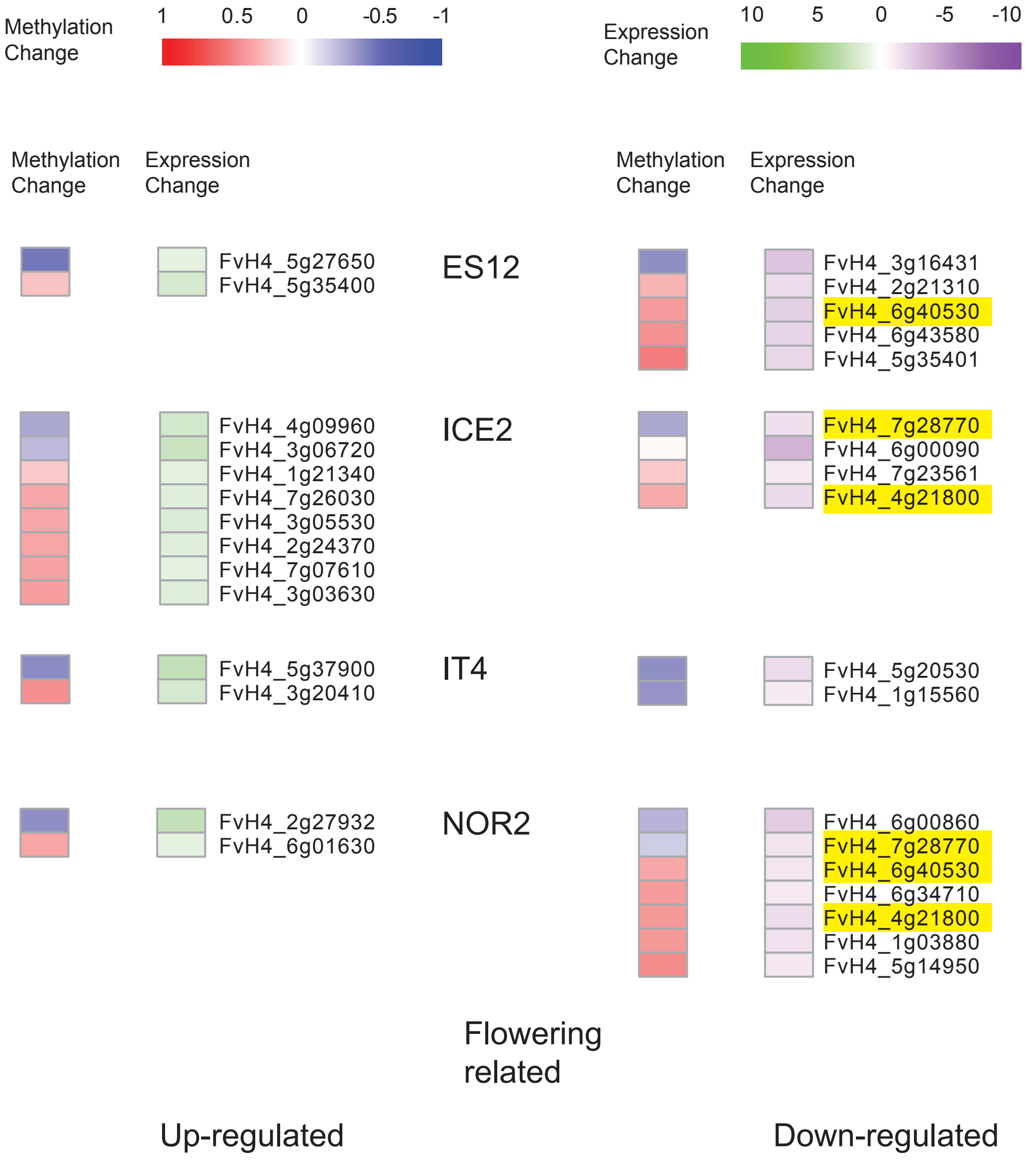

### Supplemental Figure 21

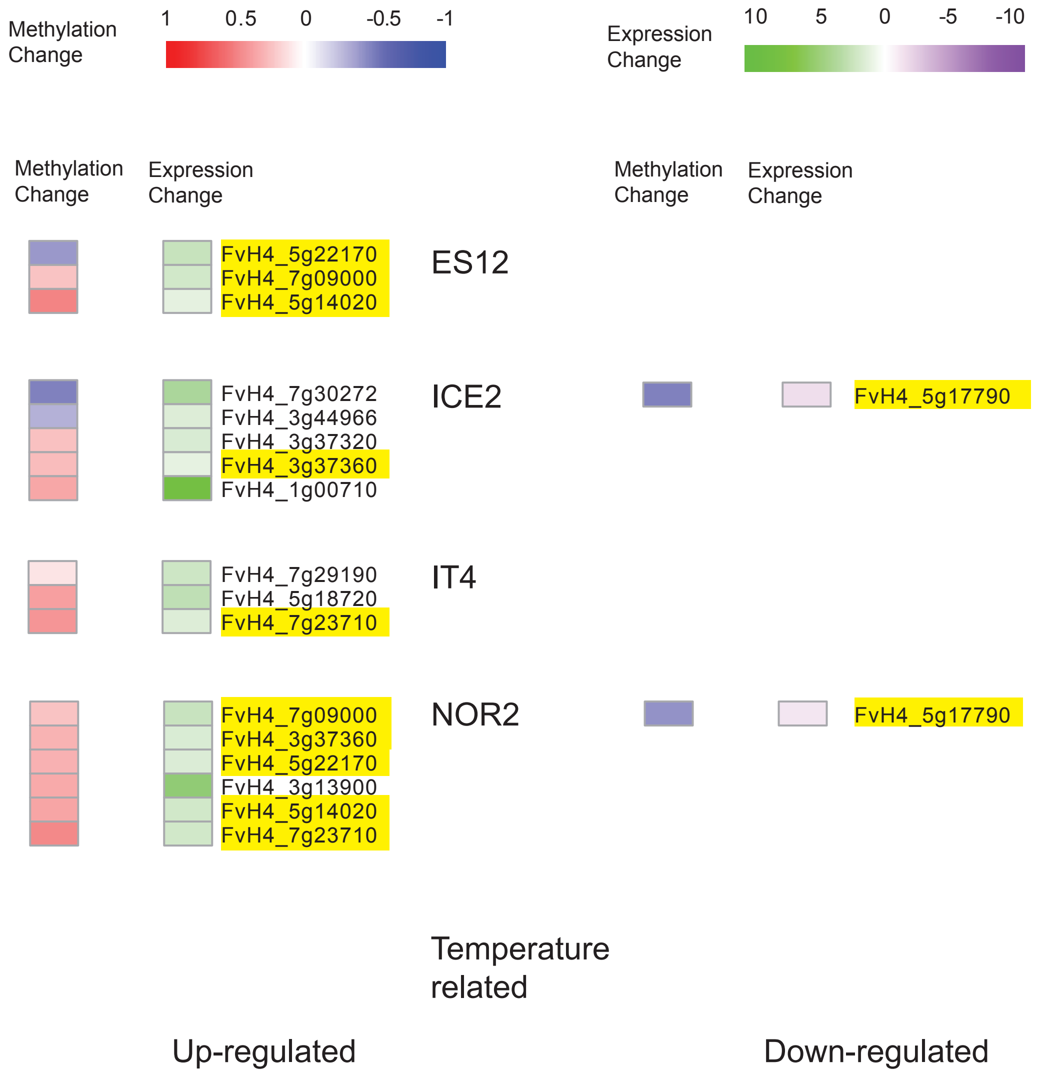

### Supplemental Figure 22

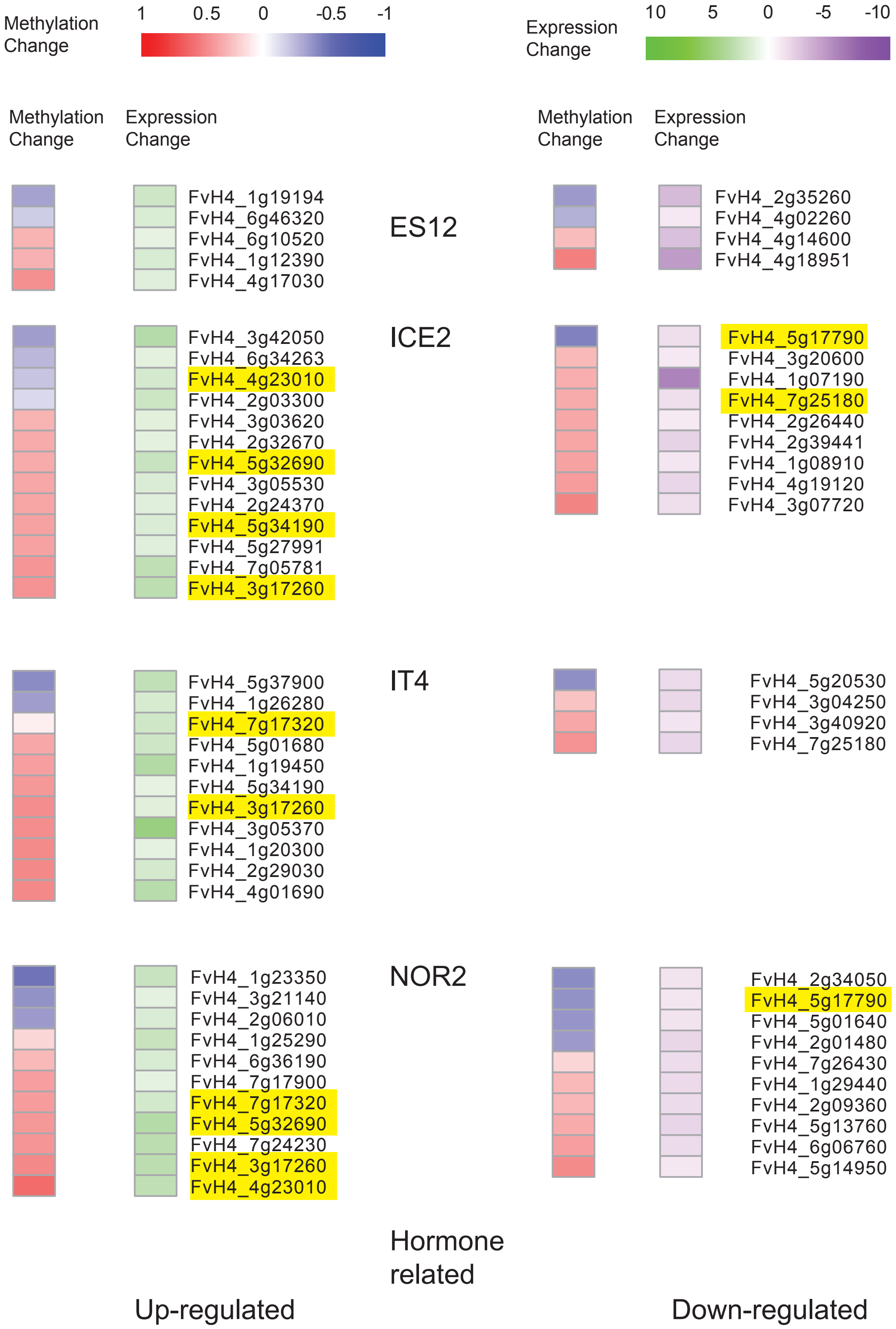

### Supplemental Figure 23

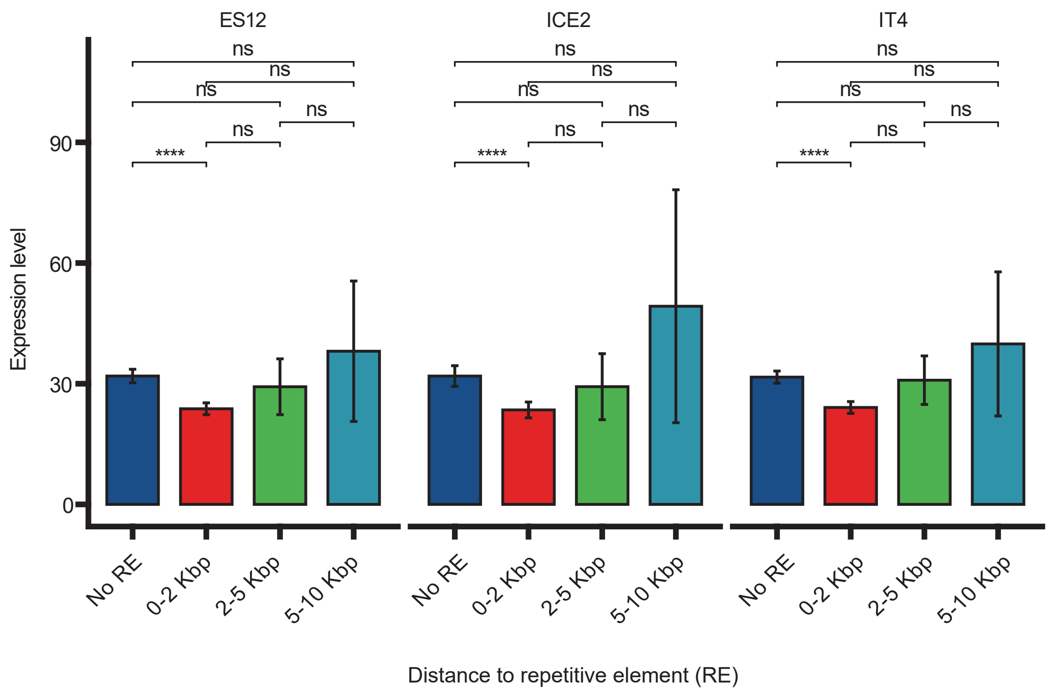

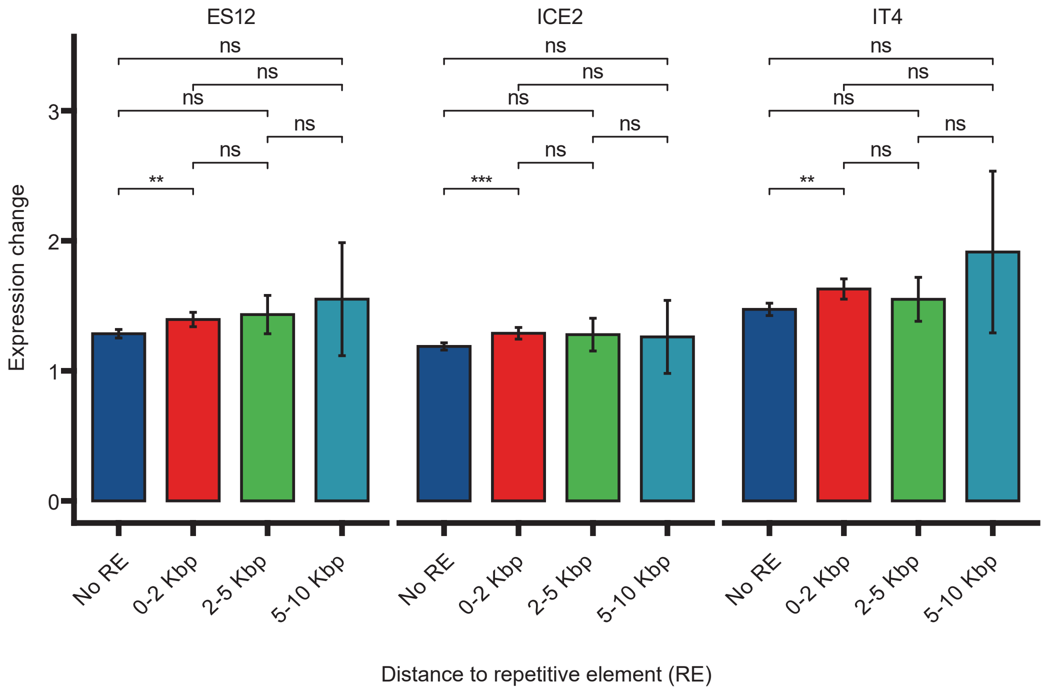
