## Supplemental Figure 17 for "Temperature-induced methylome changes during asexual reproduction trigger transcriptomic and phenotypic changes in *Fragaria vesca*"

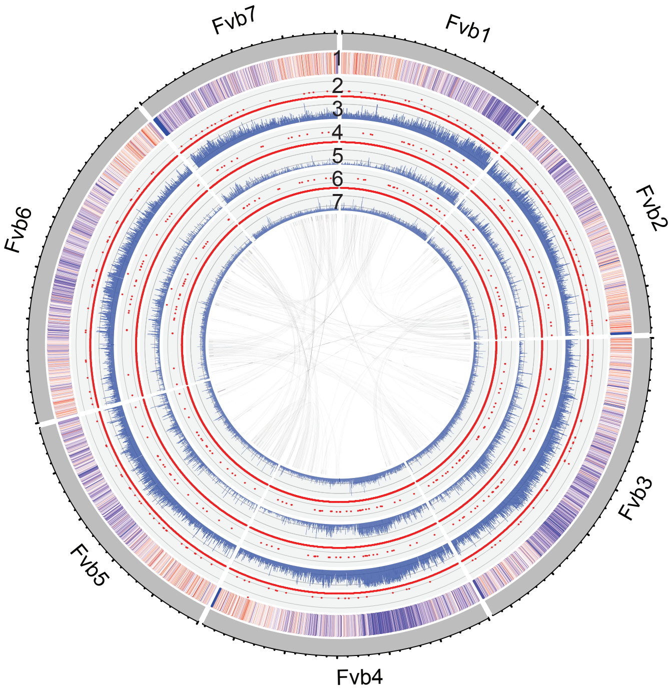


ES12

1: Gene density
2: CGN DMRs
3: CGN methylation
4: CHG DMRs
5: CHG methylation
6: CHH DMRs
7: CHH methylation

Gene Density:
Low High

Scale:
6Mbp

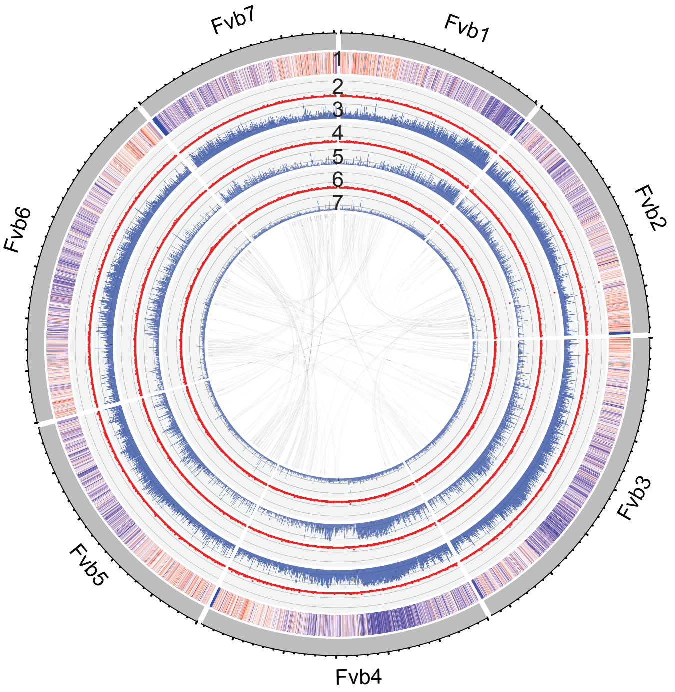


ICE2

1: Gene density
2: CGN DMRs
3: CGN methylation
4: CHG DMRs
5: CHG methylation
6: CHH DMRs
7: CHH methylation

Gene Density:
Low High

Scale:
6Mbp

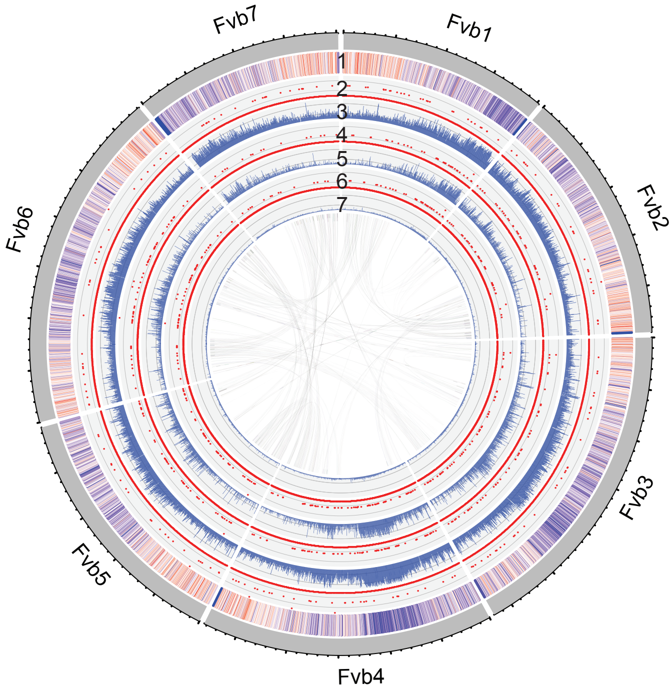


IT4

1: Gene density
2: CGN DMRs
3: CGN methylation
4: CHG DMRs
5: CHG methylation
6: CHH DMRs
7: CHH methylation

Gene Density:
Low High

Scale:
6Mbp

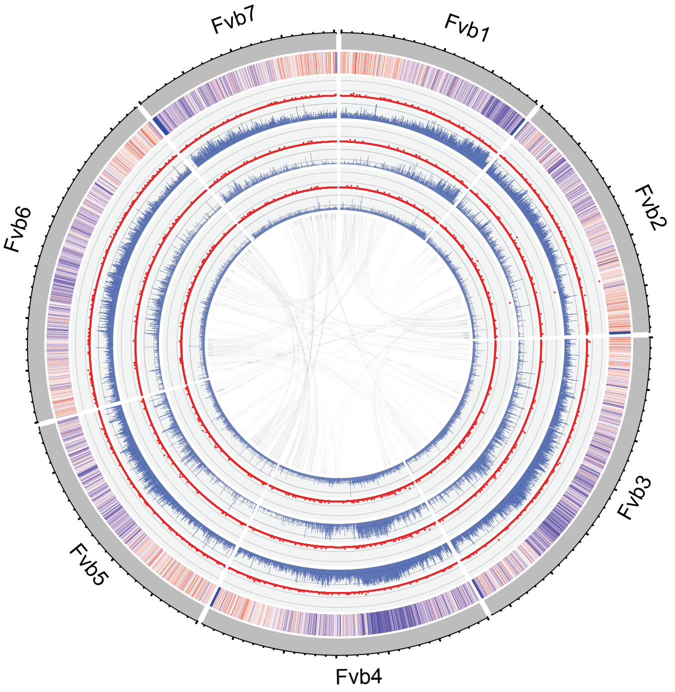


NOR2

1: Gene density
2: CGN DMRs
3: CGN methylation
4: CHG DMRs
5: CHG methylation
6: CHH DMRs
7: CHH methylation

Gene Density:
Low High

Scale:
6Mbp
